# SILAC-based quantification reveals modulation of the immunopeptidome in BRAF and MEK inhibitor sensitive and resistant tumor cells

**DOI:** 10.1101/2024.08.08.606999

**Authors:** Melissa Bernhardt, Anne Rech, Marion Berthold, Melina Lappe, Jan-Niklas Herbel, Florian Erhard, Anette Paschen, Bastian Schilling, Andreas Schlosser

**Affiliations:** Rudolf Virchow Center, Center for Integrative and Translational Bioimaging, Julius-Maximilians-Universität of Würzburg, Würzburg, Germany; Department of Dermatology, Venereology and Allergology, University Hospital Würzburg, Würzburg, Germany; Institute for Pharmacology and Toxicology, Julius-Maximilians-Universität of Würzburg, Würzburg, Germany; Faculty for Informatics and Data Science, University of Regensburg, Regensburg, Germany; Department of Dermatology, University Hospital Essen, University Duisburg-Essen and German Cancer Consortium (DKTK); Department of Dermatology, Venerology and Allergology, Goethe University Frankfurt, University Hospital, Frankfurt, Germany

**Keywords:** mass spectrometry, stable isotope labeling, HLA-I peptides, T-cell epitopes, de novo peptide sequencing, cryptic HLA peptides, melanoma, MAPK pathway inhibition

## Abstract

The immunopeptidome is constantly monitored by T cells to detect foreign or aberrant HLA peptides. It is highly dynamic and reflects the current cellular state, enabling the immune system to recognize abnormal cellular conditions, such as those present in cancer cells. To precisely determine how changes in cellular processes, such as those induced by drug treatment, affect the immunopeptidome, quantitative immunopeptidomics approaches are essential. To meet this need, we developed a pulsed SILAC-based method for quantitative immunopeptidomics. Metabolic labeling with lysine, arginine, and leucine enabled isotopic labeling of nearly all HLA peptides across all allotypes (> 90% on average). We established a data analysis workflow that integrates the *de novo* sequencing-based tool Peptide-PRISM for comprehensive HLA peptide identification with MaxQuant for accurate quantification. We employed this strategy to explore the modulation of the immunopeptidome upon MAPK pathway inhibition (MAPKi) and to investigate alterations associated with early cellular responses to inhibitor treatment and acquired resistance to MAPKi. Our analyses demonstrated significant changes in the immunopeptidome early during MAPKi treatment and in the resistant state. Moreover, we identified putative tumor-specific cryptic HLA peptides linked to these processes that might represent exploitable targets for cancer immunotherapy.

**Synopsis:** We have developed a mass spectrometric method that allowed us to investigate the effects of common MAPK inhibitors on the immunopeptidome of melanoma cells. This approach has led to the discovery of new potential targets for cancer immunotherapy.

## Introduction

The immune surveillance for self and non-self relies significantly on the processing and presentation of antigens as short peptides by major histocompatibility complex (MHC) molecules on the cell surface, enabling T cells to recognize them. MHC class I molecules are expressed in most nucleated cells and primarily present endogenously derived peptides to CD8+ T cells (1). The collection of these peptides is termed the immunopeptidome, which enables T cells to distinguish normal from abnormal and to eliminate infected as well as malignant cells. The concept of utilizing HLA peptides as targets for immunotherapeutic intervention is an active area of research, particularly in the context of anti-viral therapy, autoimmune disease and cancer immunotherapy. In the field of cancer immunotherapy, the immunopeptidome is crucial for identifying tumor-specific antigens that can be recognized by T cells, thereby driving the development of immunotherapies applying for instance T cell receptor–transduced T cells or patient-specific mRNA or peptide vaccines (2).

The pharmacological inhibition of the mitogen-activated protein kinase (MAPK) signaling pathway by inhibitors of the BRAF and MEK kinases (BRAFi and MEKi) plays an important role in the treatment of patients with advanced malignant melanoma. The sustained activation of the MAPK signaling pathway shows a strong correlation with the molecular pathogenesis of malignant melanoma (3). Dysregulation of this pathway, often due to activating mutations, results in increased signaling activity facilitating cell proliferation, invasion, metastasis, migration, survival, and angiogenesis (4). The most common mutation in this context, affecting approximately half of melanoma patients, is found in the *BRAF* gene, predominantly at codon 600 (V600E) (3). The constitutive activation of BRAF is inhibited by specific BRAFi, such as vemurafenib, which is routinely combined with inhibitors against its downstream target MEK. These MEKi, such as cobimetinib or trametinib, are used to enhance therapy efficacy (5). It has been shown that MEKi alter the immunopeptidome by upregulating tumor associated antigens (TAAs) (6). However, effects of MAPK signaling inhibition on cryptic peptides that originate from non-coding regions has not been evaluated so far.

Targeted therapies have improved progression free survival of advanced BRAF^V600^-mutated melanoma patients, but acquired resistance occurs in 50% of patients within one year after initiation of therapy (7). Initially, tumors respond to therapy, followed by a relapse of tumor cell growth upon enduring kinase inhibitor exposition. This phenomenon is prominent in many other targeted therapies involving kinase inhibitors (8). There are multiple resistance mechanisms, most of which lead to the reactivation of ERK even in the presence of inhibitors, e.g. by BRAF amplifications or by upregulation of receptor tyrosine kinases that activate alternative survival pathways such as PI3K/AKT signaling (9). Therapy of drug-resistant tumors is a significant challenge in oncology. The effects of ongoing resistance mechanisms on the immunopeptidome can be critical for guiding therapy decisions, such as indicating immunotherapies for late-stage kinase inhibitor resistant cells. We therefore aimed to study how acquired BRAF and MEK kinase inhibitor resistance directly shapes the immunopeptidome.

Mass spectrometry (MS) has proven as a powerful high-throughput method for the direct identification and characterization of the immunopeptidome (10). In recent years, there has been a substantial increase in the number of immunopeptidome datasets. These datasets encompass a wide range of samples, such as cell lines, plasma samples, different tumor entities, as well as healthy tissues. This led to the establishment of comprehensive peptide ligand atlases, such as the caAtlas (11), the HLA Ligand Atlas (12) or the Immune Epitope Database (IEDB) (13). Optimization of sample preparation (14, 15) and more sensitive instrumentation (16) recently led to improved immunopeptidome coverage. On the other hand, compared to other areas in proteomics, application of quantitative approaches is still lacking behind in immunopeptidomics. Quantitative MS-based methods, such as label-free quantification (LFQ) (17), isobaric labeling (TMT labeling) (18) and metabolic labeling (SILAC) (19) have been applied only rarely in immunopeptidomics so far.

Studying changes of the immunopeptidome in the acute phase of a cellular perturbation, would be highly valuable to improve our understanding of HLA peptide presentation, as well as for the identification of more specific and more effective T-cell targets for immunotherapy. However, accurate quantitative analysis of immunopeptidomes is hampered by several additional challenges: (i) the immunopeptidome is highly dynamic and can be strongly modulated even by subtle variations of the cellular status, (ii) peptide-level quantification is in general less accurate and more error-prone compared to protein-level quantification, (iii) reproducibility of MHC peptide isolation methods is limited, and finally, (iv) low peptide intensities and low immunopeptidome coverage lead to a high number of missing values. Altogether, this calls for a highly accurate quantification method. In contrast to LFQ and isobaric-labeling, SILAC allows to pool differentially treated cells before the affinity-based isolation of MHC peptides, thereby greatly improving accuracy for relative quantification. Thus, we established an optimized SILAC-based workflow that is customized for immunopeptidomics and enables reproducible and accurate quantification of HLA-I peptides. We validated this method by inducing interferon gamma (IFNγ)-dependent HLA-I peptide alterations (20) and applied this workflow to reveal quantitative changes in the immunopeptidome of melanoma cell lines upon treatment with the BRAFi/MEKi combination vemurafenib and cobimetinib and the MEKi trametinib alone and compared the immunopeptidomes of a resistant and a non-resistant variant of a melanoma cell line.

## Materials and Methods

### Cell Culture and Stable Isotope Labeling with Amino Acids (SILAC)

Melanoma cell lines Ma-Mel-63a (HLA-A*02:01,HLA-A*03:01,HLA-B*07:02,HLA-B*18:01,HLA-C06:02, provided by Dirk Schadendorf, University Hospital Essen, Germany), SK-Mel-28 (HLA-A*11:01, HLA-B*40:01, HLA-C*03:03), UKE-Mel-105b (HLA-A*03:01, HLA-B*25:01, HLA-C*04:01, University Hospital Essen, Germany) were cultured at 37°C with 5% CO_2_ under humidified atmosphere. Ma-Mel-63a and SK-Mel-28 were grown in RPMI1640 medium (Thermo Fisher) supplemented with 10% FCS (Capricorn) and 1% penicillin/streptomycin (P/S, Sigma-Aldrich). UKE-Mel-105b cells were cultured in DMEM high glucose HEPES medium (Gibco) supplemented with 10% FCS and 1% P/S. A SK-Mel-28 vemurafenib and cobimetinib double resistant cell line was established. As the dual treatment with both inhibitors is challenging for most melanoma cell lines, first the resistance to vemurafenib (v) was introduced. Subsequently, the v-resistant cell lines underwent treatment with cobimetinib (c), to establish double resistant cell lines (SK-Mel-28 rr). For this purpose, cells were initially cultured in a low concentration of 15 nM vemurafenib (PLX4032, Cayman Chemical) in RPMI-CM (RPMI-1640 Medium with L-glutamine and sodium bicarbonate, Sigma Aldrich, + 10 % FBS superior, Biochrom + 1% P/S). The medium was refreshed every 2-3 days, and regular detachment of cells was performed to prevent excessive adherence. Upon reaching 80-90% confluence, the cells were split at a ratio of 1:4 and allowed to grow again to 80-90% confluence under the same inhibitor concentration. The doubling of the inhibitor concentration occurred only during the second 1:4 split. This iterative process continued until the cells adapted to grow in 2 µM Vemurafenib. Subsequently, a portion of the cells was cryopreserved, while the remaining cells were subjected to the same regimen with 2 µM vemurafenib and an additional 1.5 nM cobimetinib (GDC-0973, Cayman Chemical) until a final concentration of 200 nM cobimetinib was attained. The resistance profiles of the 2 µM v-resistant lines (r) and the 2 µM v- and 0.2 µM c-resistant lines (rr) were then evaluated through Western blot analysis.

Metabolic labeling was conducted using heavy (H) and medium-heavy (M) isotopically labeled amino acids arginine-13C6-15N4 (Arg10), lysine-13C6-15N2 (Lys8), leucine-13C6 (Leu6) or leucine-13C6-15N1 (Leu7) and arginine-13C6 (Arg6), lysine-D4 (Lys4), leucine-D3 (Leu3), respectively. All isotopically labeled amino acids were purchased from Eurisotop. RPMI-1640 for SILAC was supplemented with 90 mg/L isotopically labeled arginine, 40 mg/L isotopically labeled lysine and optionally 50 mg/L isotopically labeled leucine. L-Proline was added in a concentration of 180 mg/L to suppress arginine to proline conversion. DMEM for SILAC was supplemented with 1 M HEPES (cell culture grade, Sigma Aldrich) to a final concentration of 24 mM, followed by 69 mg/L isotopically labeled arginine, 117 mg/L isotopically labeled lysine and 105 mg/L isotopically labeled leucine. Additionally, 10% dialyzed FCS (Capricorn) and 1% P/S (Sigma Aldrich) was added to both RPMI-1640 and DMEM SILAC media. Cells were either expanded in SILAC medium for more than five cell division cycles, resulting in complete labeling, or labeled over a defined time interval with SILAC medium (pulsed SILAC, pSILAC) while a treatment is applied. SK-Mel-28 wildtype (wt) cells were grown in RPMI SILAC medium containing medium-heavy labeled amino acids (RPMI M). SK-Mel-28 vemurafenib/cobimetinib double resistant cells (SK-Mel-28 rr) were grown in RPMI SILAC medium supplemented with heavy labeled amino acids (RPMI H). Ma-Mel-63a cells were grown in parallel in RPMI M and RPMI H, respectively. The following pSILAC experiments were conducted: Ma-Mel-63a cells were treated for 24h and 72h with 100 IU IFN-gamma (IFNγ, Peprotech) using RPMI M. In parallel, a vehicle control experiment with H_2_O was conducted for 24h and 72h using RPMI SILAC H. Ma-Mel-63a cells were further treated with (i) a combination of 1 µM vemurafenib plus 100 nM combimetinib for 4h, 8h, 21h, 48h and (ii) 100 nM trametinib (MedChem Express) for 24h and 72h. The treatment was performed using RPMI H, while the vehicle control experiment was conducted in RPMI M containing 0.1% DMSO. Pulsed-SILAC experiments, including a vemurafenib + cobimetinib treatment and a control experiment were performed in the same manner for SK-Mel-28 wt cells and SK-Mel-28 rr for 24h and 72h. UKE-Mel-105b cells were treated with 100nM trametinib for 24h and 72h using DMEM for SILAC supplemented with heavy-isotope labeled amino acids (DMEM H) and DMEM with medium-heavy isotope labeled amino acids (DMEM M) for the corresponding vehicle control experiment (0.1% DMSO). For each cell line, an additional control experiment with mock treatment for both labeling conditions was performed. Cells were harvested, aliquoted at equal cell numbers and stored at -80°C until immunopeptidome analysis. For each SILAC condition 2-5x 10^7^ cells were used.

### Primary Melanocytes

Normal human epidermal melanocytes, M2 (NHEM) from a 5-year-old, male, Caucasian healthy donor (HLA-A*03:01, HLA-A*68:01, HLA-B*07:02, HLA-B*44:02. HLA-C*07:02, HLA-C*07:04) were purchased from PromoCell and cultivated in Melanocyte Growth Medium M2 (PromoCell) until 90% confluency was reached. Cells were harvested for qualitative immunopeptidome analysis.

### Immunoaffinity Purification and Isolation of HLA-I Peptides

Cell pellets were lysed in 2 ml lysis buffer per 1x 10^8^ cells for 1h at 4°C. Cell lysates were centrifuged at 16,000 x g for 20min at 4°C for preclearing. Precleared lysates of heavy-isotope labeled cells were combined in a 1:1 ratio with medium-heavy isotopically labeled cells, resulting in a mixture of treatment and control cells. Immunoaffinity purification, HLA-I peptide isolation were conducted as described previously using W6/32 antibody (kindly provided by Hans-Georg Rammensee, Department of Immunology, University of Tuebingen, Germany) non-covalently linked to Sepharose beads, followed by solid phase extraction (SPE) with restricted access material (RAM) (15). Eluted HLA-I peptides were lyophilized and stored at -20°C until LC-MS/MS analysis.

### Proteome Analysis

5 µl of cell lysate was precipitated with the fourfold volume of acetone overnight at -20 °C. Pellets were washed with acetone at -20 °C. Precipitated proteins were dissolved in NuPAGE® LDS sample buffer (Life Technologies), reduced with 50 mM DTT at 70 °C for 10 min and alkylated with 120 mM iodoacetamide at room temperature for 20 min. Separation was performed on NuPAGE® Novex® 4-12 % Bis-Tris gels (Life Technologies) with MOPS buffer according to manufacturer’s instructions. The gel was washed three times for 5 min with water and stained for 1 h with Simply Blue™ Safe Stain (Life Technologies). After washing with water for 1 h, each gel lane was cut into 15 slices. The excised gel bands were destained with 30 % acetonitrile in 0.1 M NH_4_HCO_3_ (pH 8), shrunk with 100 % acetonitrile, and dried in a vacuum concentrator (Concentrator 5301, Eppendorf, Germany). Digests were performed with 0.1 µg trypsin (Trypsin Gold, Mass Spectrometry Grade, Promega) per gel band overnight at 37 °C in 0.1 M NH_4_HCO_3_ (pH 8). After removing the supernatant, peptides were extracted from the gel slices with 5 % formic acid. Extracted peptides were pooled with the supernatant and analyzed by LC-MS/MS.

### Quantitative Immunopeptidome Analysis

LC-MS/MS analysis of HLA-I peptides was performed as described previously (15). In brief, HLA-I peptides were dissolved in 30 μL of 2% ACN, 0.1% formic acid, and two subsequent LC-MS/MS runs were performed on an Orbitrap Fusion (Thermo Scientific) using a method for doubly charged peptides and a method for triple and single charged peptides. MS1 spectra were acquired with the following parameters: AGC target: 50,000, dynamic exclusion: 1, exclusion duration: 8s. MS2 spectra were acquired with the following paramters: intensity threshold: 10,000, collision energy: 30;35;40%, resolution: 60,000, AGC target: 50,000. ETD was applied for triply charged peptides only. MS data was analyzed by *de novo* sequencing with PEAKS XPro (Bioinformatics Solutions Inc.). Raw data refinement was performed with the following settings: (i) Merge Options: no merge; (ii) Precursor Options: corrected; (iii) Charge Options: 1−6; (iv) Filter Options: no filter; (v) Process: true; (vi) Default: true; (vii) Associate Chimera: yes. Parent Mass Error Tolerance was set to 10 ppm and Fragment Mass Error Tolerance was set to 0.02 Da. Enzyme was set to none.

To consider the medium-heavy and heavy labeled amino acids the following variable posttranslational modifications (PTMs) were allowed: Arg10 (Δm= 10.0083), Arg6(Δm= 6.0201), Lys8 (Δm= 8.0142), Lys4 (Δm= 4.0251), Leu6 (Δm= 6.0201) or Leu7(Δm= 7.0201), Leu3 (Δm= 3.0188). Additionally, Oxidation (M), pyro-Glu from Q (N-term Q), and carbamidomethylation (C) were included as variable PTMs. A maximum of 6 variable PTMs were allowed per peptide. Up to 10 *de novo* sequencing candidates were reported for each identified fragment ion mass spectrum, with their corresponding average local confidence (ALC) score. All *de novo* sequence candidates were matched against the 6-frame translated genome (GRCh38) and the 3-frame translated transcriptome (ENSEMBL release 90) using Peptide-PRISM (21). Peptides were then assigned to categories based on their genomic locations where they are encoded: (i) CDS, known protein; (ii) 5′-UTR; (iii) Off-frame, encoded within a protein coding region, but shifted by ±1; 3′-UTR; (iv) ncRNA; (v) Intronic; (vi) Intergenic; (vii) CDSintoIntron; (viii) OthersIntoIntron. Results were filtered to category-specific false discovery rate (FDR). NetMHCpan 4.0 was used to predict binding affinities for all identified MHC-I peptides. As per default, a cut-off of 0.5% rank for strong binders and 2% rank for weak binders was used. From all identified HLA-I peptides identified with Peptide-PRISM, a non-redundant custom fasta database was generated (category-and length-specific FDR < 10%). For each experiment, typically comprising one treatment and one mock condition for two different timepoints, an individual database was generated. After obtaining fasta databases, MaxQuant 2.4.2 (22) was used for quantitative analysis. Digestion mode was set to “no digestion”. Multiplicity for the SILAC-labeling was set to 3 (light, medium & heavy) with a maximum of 6 labels per peptide. Arg6, Lys4, Leu3 were selected as medium labels and Arg10, Lys8, Leu6 or Leu7 as heavy labels. FDR filtering was turned off by setting PSM FDR, protein FDR and site decoy fraction to 1. Minimum scores for modified and for unmodified peptides were set to 25. The second peptide option was enabled. Finally, the re-quantify option was used for improving quantification of large ratios. Apart from these adapted settings, MaxQuant default parameters were used. The MaxQuant peptides table was merged with the Peptide-PRISM results table by peptide sequence, and MaxQuant H/M ratios were used to generate scatter plots showing the modulation of the immunopeptidome. For improved ratio calculation, peptide H/M ratios were re-calculated from MaxQuant evidence table as follows (**Supplemental Table 1**): for each experimental condition (“experiment type”, “timepoint”) the “median lg2 H/M ratio” for each peptide sequence was calculated from MaxQuant H/M ratios in the evidence table. When a peptide sequence feature was detected in both, medium and heavy labeling state (“Labeling State over Raw File” = 1,2 or 0,1,2), but was not recognized as a labeling cluster (due to large retention time shift of the differentially labeled species) the corresponding H/M ratios were re-calculated from the “summed Intensity H over Experiment” and the “summed Intensity M over Experiment” (Fig. 2 and Suppl. Fig. 6) and used instead the original MaxQuant H/M ratios for calculating “median lg2 H/M ratio corrected”. To correct for potential mixing errors, “median lg2 H/M ratio corrected” was normalized on the median of the distribution. The resulting values in the columns “norm median lg2 H/M ratio corrected experiment type, timepoint” were used to generate the scatter plots in Fig. 2D, Fig. 3, Fig 5 B, D and E, Supplemental Fig. 8,10,11,12,16,17 and 18. Intensity values (“log10 (intensity)”) used in the corresponding scatter plots were calculated from the summed intensities of all labeling states over one experimental condition.

**Figure 1.**
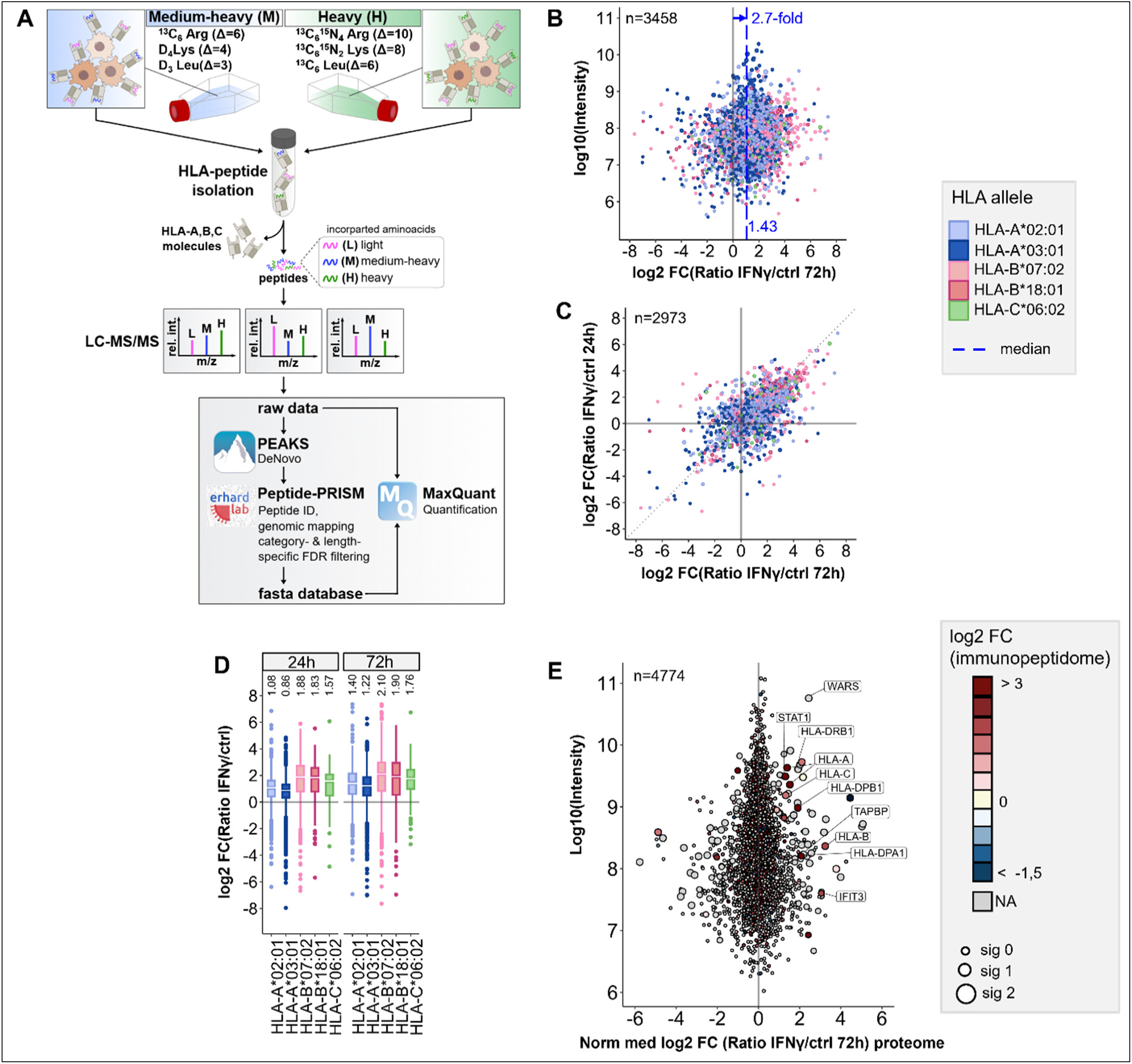
Workflow for pSILAC-based quantitative immunopeptidomics and its application to IFNγ tretment. (**A**) Workflow outline for the optimized pSILAC-based quantitative immunopeptidomics approach. Cells, initially cultured in medium with light amino acids, are transferred to medium containing either medium-heavy (M) or heavy (H) labeled amino acids. One culture undergoes a treatment, while the other serves as a control. The isotopically labeled amino acids are incorporated into HLA peptides during the labeling pulse. Combined in a 1:1 ratio, the HLA peptides from control sample and treatment sample are isolated together and subjected to LC-MS/MS analysis. *De novo* peptide sequencing is performed with PEAKS XPro, followed by HLA peptide identification, mapping to genomic locations and stratified FDR filtering using Peptide-PRISM. NetMHCpan 4.0 is applied to predict binding of the identified HLA peptides to the corresponding alleles of the cells. An experiment-specific fasta database containing all identified HLA peptides is then used for quantitative analysis with MaxQuant. (**B**) Quantitative changes of HLA-peptide presentation upon IFNγ treatment are represented by Log2-fold-change (Lg2fc) values of H/M ratios. The scatter plot displays alterations of every single HLA peptide. HLA alleles are color-coded according to their assigned HLA allele. Global peptide presentation is enhanced by a ∼2.7-fold (Lg2fc 1.43). (**C**) Correlation of quantitative Lg2fc values upon 24h and 72h IFNγ treatment. (**D**) Boxplots showing global allomorph-dependent changes in peptide presentation for both time points. (**E**) Quantitative proteomic changes are represented by normalized Lg2fc-values. Colors correspond to the median-Lg2fc values of the HLA peptides from the same protein. Immunopeptidome data was filtered by category- and length-specific FDR of <10% and included only NetMHCpan-predicted binders. Proteome data was filtered to 1% PSM-FDR and 1% protein FDR.

**Figure 2.**
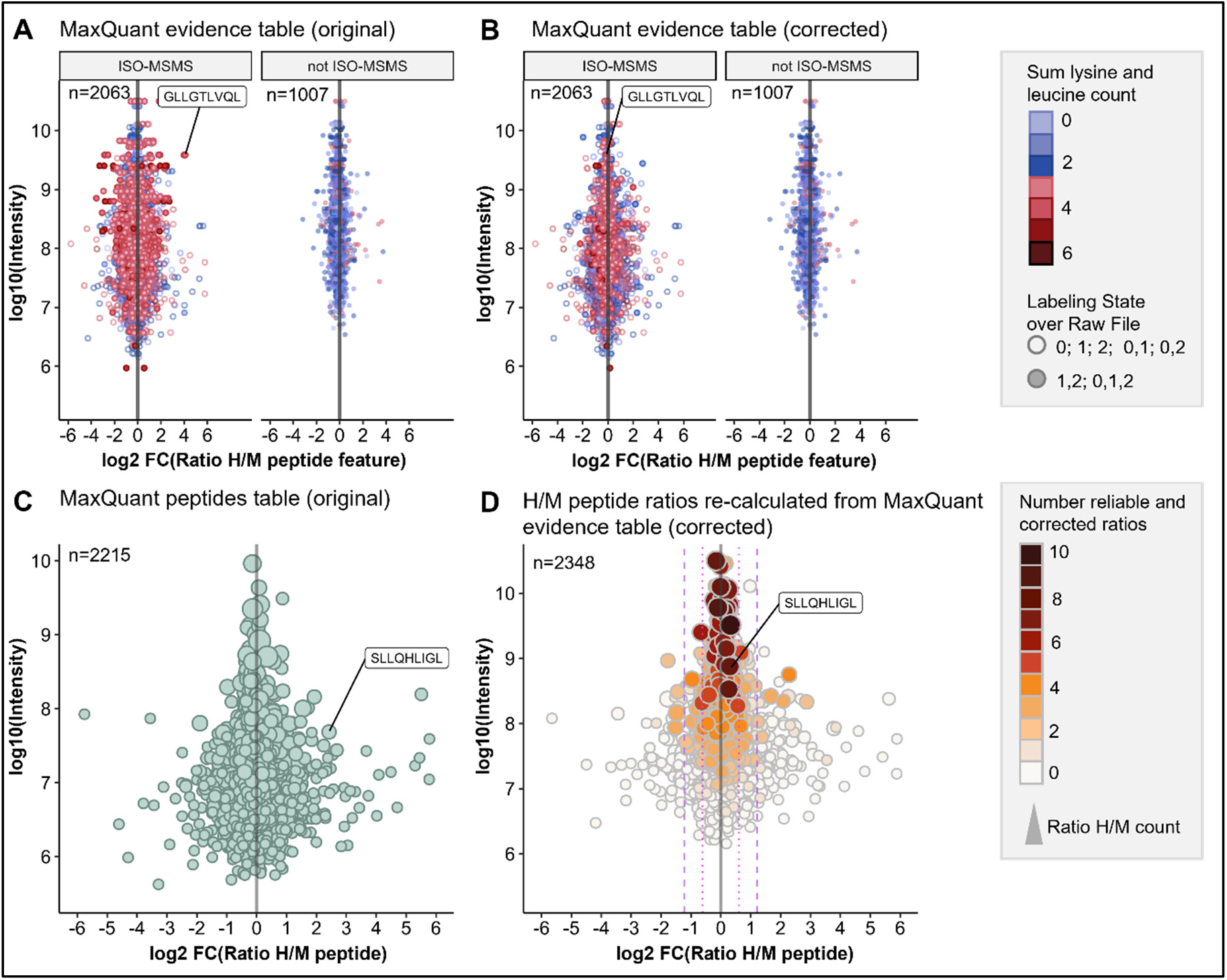
Optimized data analysis for improved quantification accuracy. pSILAC data was obtained from a 24h mock treatment experiment with Ma-Mel-63a cells. HLA peptides were identified with Peptide-PRISM (1% FDR, only NetMHCpan-predicted binders) and quantified with MaxQuant. (**A**) Original H/M ratios of all peptide features from MaxQuant evidence table for type ISO-MSMS and not ISO-MSMS (MULTI-MSMS, MULTI-MATCH-MSMS, MULTI-MATCH, MULTI-SECPEP). Colors represent the sum of lysine and leucine residues per peptide. Labeling state 0 represents light, 1 medium-heavy and 2 heavy labeled peptides. (**B**) Corrected H/M ratios of all peptide features from MaxQuant evidence table for type ISO-MSMS and not ISO-MSMS. (**C**) Original H/M peptide ratios from MaxQuant peptides table. (**D**) H/M peptide ratios re-calculated from MaxQuant evidence table (corrected). The number of reliable and corrected H/M ratios is color-coded and used as a measure for quantification accuracy.

**Figure 3.**
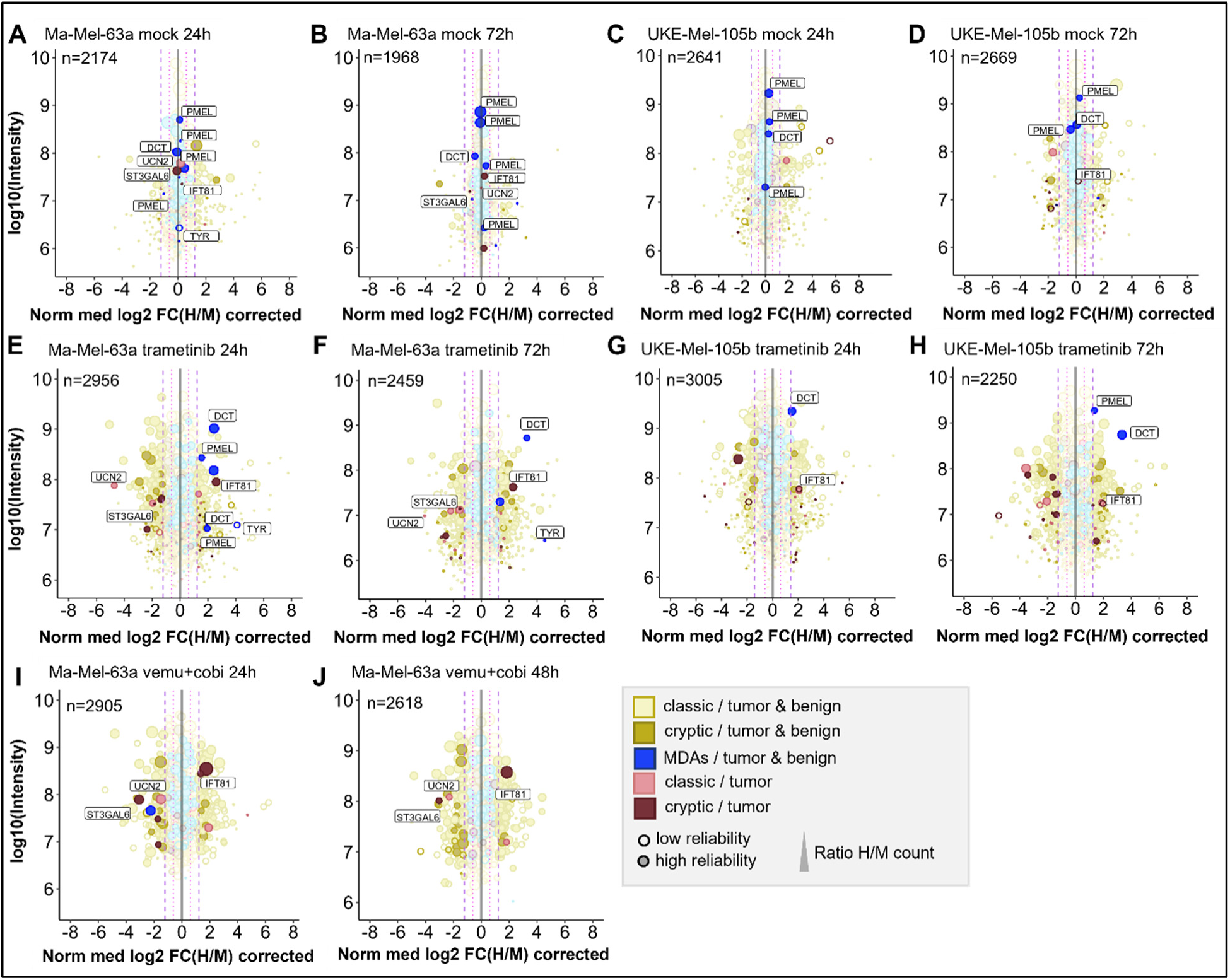
Quantitative changes of the immunopeptide repertoires of Ma-Mel-63a and UKE-Mel-105b cells upon inhibition of MAPK signaling with trametinib or vemurafenib/cobimetinib. Mock experiments were performed to estimate cell line-specific background changes occurring during labeling intervals without treatment. (A) Mock experiment with Ma-Mel-63a after 24h. (B) Mock experiment with Ma-Mel-63a after 72h. (C) Mock experiment with UKE-Mel-105b after 24h. (D) Mock experiments with UKE-Mel-105b after 72h. Peptides that are regulated upon BRAF inhibition or MEK inhibition are also highlighted in mock conditions. (E) Ma-Mel-63a cells were treated with 100 nM trametinib for 24h. (F) Ma-Mel-63a cells were treated with 100 nM trametinib for 72h. (G) UKE-Mel-105b cells were treated with 100 nM trametinib for 24h. (H) UKE-Mel-105b cells were treated with 100 nM trametinib for 72h. (I) Ma-Mel-63a cells were treated with 1 µM vemurafenib and 100 nM trametinib for 24h. (J) Ma-Mel-63a cells were treated with 1 µM vemurafenib and 100 nM trametinib for 48h. Immunopeptidomes were globally altered upon treatment. Presentation of MDAs was induced. The presentation of a putative tumor-exclusive peptide RVYLDIVTPK *(IFT81)* was induced upon treatment, while presentation of the peptide RLSSSLPSR (*ST3GAL6*) was diminished. MDA = melanocyte differentiation antigens. CGAs = cancer/germline antigens.

For quantification quality assessment H/M ratios of type “multi” were classified as “reliable” in the column “Ratio reliability”, H/M ratios of type “iso” were classified as “unreliable”, except those that were detected in both, medium and heavy labeling state (“Labeling State over Raw File” = 1,2 or 0,1,2), but not recognized as a labeling cluster. These were corrected as described above and classified as “corrected”. The values from the column “Number of reliable and corrected ratios” can be used to color-code highly reliable and less reliable H/M ratios (**Figure 2D**).

### Tumor-versus-Healthy Database (TvHdb)

Publicly available immunopeptidome datasets of tumor and healthy cells (**Supplemental Table 2**) were re-analyzed by *de novo* sequencing with PEAKS XPro and Peptide-PRISM as described above. We analyzed ∼2500 MS raw data files from ∼350 different patients/cell lines and identified more than 300,000 unique HLA-I peptides (<5% false discovery rate (FDR), ALC>50, only NetMHCpan-predicted binder), presented on ∼140 different HLA allomorphs. Tumor data comprises samples from melanoma, meningioma, acute myeloid leukemia, colorectal cancer, ovarian cancer, glioblastoma and several other entities. Healthy control data includes the complete data set of the HLA ligand atlas project (12) (hla-ligand-atlas.org) comprising autopsy samples from 29 tissues from 13 donors, a large variety of samples from healthy blood donors, and some matched healthy tissue samples. We mapped all identified peptides with a unique genomic location to the human genome (hg38) and to their corresponding open reading frames (ORFs) and classified those ORFs as tumor-exclusive from which HLA peptides were detected exclusively in tumor samples.

### Quantitative Proteome Analysis

Proteome analyses were performed by nanoLC-MS/MS on an Orbitrap Fusion (Thermo Scientific) equipped with a PicoView Ion Source (New Objective) and coupled to an EASY-nLC 1000 (Thermo Scientific). Peptides were loaded on a trapping column (2 cm x 150 µm ID, PepSep) and separated on a capillary column (30 cm x 150 µm ID, PepSep) both packed with 1.9 µm C18 ReproSil and separated with a 45-minute linear gradient from 3% to 30% acetonitrile and 0.1% formic acid and a flow rate of 500 nl/min. Both MS and MS/MS scans were acquired in the Orbitrap analyzer with a resolution of 60,000 for MS scans and 30,000 for MS/MS scans. HCD fragmentation with 35 % normalized collision energy was applied. A Top Speed data-dependent MS/MS method with a fixed cycle time of 3 s was used. Dynamic exclusion was applied with a repeat count of 1 and an exclusion duration of 30s; singly charged precursors were excluded from selection. Minimum signal threshold for precursor selection was set to 50,000. Automatic gain control (AGC) was used with manufacturer’s standard settings for MS scans and MS/MS scans. EASY-IC was used for internal calibration. Raw MS data files were analyzed with MaxQuant version 2.0 (22). The search was performed against the UniProt reference proteome database (April 2024,104581 entries) with tryptic cleavage specificity, allowing 3 missed cleavages. Protein identification was under control of the false discovery rate (FDR; 1% FDR on protein and peptide spectrum match (PSM)). In addition to MaxQuant default settings, the search was performed against following variable modifications: Protein N-terminal acetylation, peptide N-terminal pyroglutamate formation at glutamine and oxidation (Met). Carbamidomethyl (Cys) was set as fixed modification. Arg6, Lys4 and Leu3 were set for medium-heavy SILAC labels and Arg10, Lys8 and Leu6 or Leu7 for heavy SILAC labels. Further data analysis was performed using R scripts developed in-house. For quantification of pSILAC-labeled proteins, the median was calculated from log2-transformed normalized peptide heavy-to-medium ratios (H/M) for each protein. Two ratio counts were required for protein quantification.

### Western Blotting

Protein extraction utilized RIPA lysis buffer from frozen dry pellets. Total protein concentration was determined via the BCA protein assay (Thermo Fisher). Equal protein quantities (12 µg per lane for 24h/48h/72h samples and 20 µg per lane for resistance validation) were combined with Pierce Lane Marker Reducing Sample Buffer (Thermo) and applied onto SDS gels (Mini-PROTEAN TGX Gels 8-16%, BioRad for 24h/48 h/72h samples and a 10% SDS gel for resistance validation). Two identical gels were loaded per time point, labeled Membrane 1 (M1) and Membrane 2 (M2), The Page Ruler Plus Prestained Protein Ladder, 10-250 kDa (Thermo), served as a marker. SDS-PAGE for protein separation was conducted in 1x Electrophoresis Buffer (25 mM Tris, 192 mM Glycine, 0.1% SDS) at a constant 120 V. Transfers onto 0.45 µm Nitrocellulose Membranes (Cytiva) occurred in 1x Blotting Buffer (48 mM Tris, 39 mM Glycine, 20% Methanol, 1.24 mM SDS) at a constant 100 V for 45 min using a CriterionTM Blotter (Bio Rad). Following Ponceau S staining to verify protein transfer integrity, membranes were treated with primary antibodies (**Supplemental Table 3)**, diluted to 1:1000 in TBS-T, 5% milk containing 0.01% NaN_3_. For 24h/48h/72h samples, the application sequence was ERK followed by Vinculin or pERK, HLA-A, B, C followed by Vinculin. Resistant samples used primary antibodies pERK, Akt followed by Vinculin or pAkt, pERK followed by Vinculin. HRP-conjugated secondary antibodies (**Supplemental Table 3**) were applied at a concentration of 1:2,000 in TBS-T/5% milk, depending on the species of the primary antibody. Development occurred with an ECL solution (Enhanced Chemiluminescence, 0.1 M Tris pH 8.5, 1.25 mM Luminol, 0.198 mM p-coumaric acid, 0.009% H_2_O_2_) using an Amersham Imager 600 (GE Healthcare Life Sciences).

### *In Vitro* Priming assay

For immunogenicity testing of selected peptide candidates, the *in vitro* priming strategy described by Bozkus *et al.* (23) was used with minor modifications. Peripheral blood mononuclear cells (PBMCs) of healthy donors were isolated from leukoreduction system chambers obtained from the Institute for Transfusion Medicine and Hemotherapy (University Hospital Wuerzburg, Germany). Therefore, Histopaque®-1077(Sigma Aldrich) density gradient separation was performed. Isolated PBMCs were frozen as previously described (23) to maintain cell viability. HLA-typing was performed via HLA-surface staining with anti-HLA*02-APC-antibody (Miltenyi Biotec) and anti-HLA-A*03-FITC-antibody (Milteny Biotec), followed by flow cytometry analysis. HLA-A*02 and/or HLA-A*03 positive donors were used for *in vitro* priming assays. Frozen PBMCs were thawed and seeded in 96-well plates (Corning). PBMCs were cultured at 37°C with 5% CO_2_ under humidified atmosphere and differentiated into monocyte derived dendritic cells (moDCs) using X-Vivo 15 Serum-free Hematopoietic Cell Medium (Lonza) medium supplemented with human GM-CSF (Gentaur), human IL-4 (Bio-Techne) and Flt3-Ligand (Bio-Techne) A fresh aliquot of donor cells was thawed and the MACS CD8+ T-cell isolation kit (Miltenyi Biotec) was used according to the manufacturers instruction to enrich for CD8 positive cells. Supernatants of each well, including non-adherent PBMCs were removed and substituted by the CD8 positive enriched T-cell fraction. Subseqeuntly, cells were matured with LPS (Sigma Aldrich), R848 (MedChem Express), IL-1β (Biotechne) and stimulated with peptides in a final concentration of 2µg/ml. The expansion of T-cells was supported by adding IL-2, IL-7 (Bio-Techne) and IL-15 (PeproTech) as described. On day 9, re-stimulation of peptide was induced according to the protocol of Boskus *et al.* (23). Cells were stained with surface antibodies anti-CD3-PE-Vio, anti-CD4-APC-Vio770 and anti-CD8-FITC (Miltenyi Biotec). Fixation and permeabilization were performed using the BD Cyofix kit and BD Perm/Wash kit (Becton Dickinson). Intracellular staining was conducted using anti-TNFα-APC and anti-IFN-γ-PE antibodies (Miltenyi Biotec). A VioBlue-live-dead marker (Thermo Fisher) was included. Flow Cytometry was performed using the MACSQuant 16 Analyzer (**Supplemental Figure 1**).

## Results

### Proof of concept – Modulation of the immunopeptidome by IFNγ

To get a first impression of the applicability, accuracy and reproducibility of pulsed SILAC (pSILAC) (24) for quantitative immunopeptidomics, we performed a proof-of-concept experiment to monitor the modulation of the immunopeptidome of a melanoma cell line by IFNγ. The effects of IFNγ on the immunopeptidome are well studied and could be used to assess the outcome of this quantitative approach. To this end we treated Ma-Mel-63a cells with 100 IU/mL IFNγ or vehicle control in medium-heavy (Lys4 & Arg6) or heavy (Lys8 & Arg10) SILAC medium, respectively. The differentially labeled IFNγ-treated and mock-treated cells were pooled after 24h and 72h, respectively. HLA-I peptides were isolated and analyzed by nanoLC-MS/MS. We used PEAKS for *de novo* peptide sequencing and Peptide-PRISM to identify both conventional peptides being part of the known proteome as well as cryptic peptides encoded in mRNA sequences that are not part of the protein-coding open reading frames, from non-coding RNAs, from introns or intergenic regions with a class-specific FDR of 10% as previously described (21). Next, we used MaxQuant for the quantification of all filtered peptides (22), and finally merged the results from Peptide-PRISM and MaxQuant (peptides table) (**Figure 1A**).

We observed a 2.1-fold median global upregulation of HLA peptides by IFNγ after 24h (**Supplemental Figure 2A**) and a 2.7-fold upregulation after 72h (**Figure 1B**), together with a strong up- and downregulation of many individual peptides, as well as a good correlation of HLA peptide-specific up- and downregulation at 24 and 72h (**Figure 1C**). In addition, we recognized the most profound upregulation of peptides presented on HLA-B (median ∼4-fold) compared to peptides presented on HLA-A alleles (median ∼2-fold) (**Figure 1D**). A comparison of quantitative immunopeptidomics and proteomics data showed no significant correlation, except for some proteins of the IFNγ-signaling pathway (e.g STAT, WARS) or the HLA peptide presentation pathway, such as HLA class I and HLA class II (**Figure 1E**; **Supplemental Figure 2B**). The increased abundance of HLA peptides derived from these gene can be explained by the global IFNγ-induced expression of these genes on transcriptome and proteome level. These results are consistent with prior findings, including transcriptome studies and recent analyses of immunopeptidomics, as well as proteomics and genomics examining the effects of IFNγ (14)(25). In summary, the proof-of-concept experiment indicated that the pSILAC approach is well suited for quantitative immunopeptidomics, gives reproducible results, and enables to monitor the modulation of the immunopeptidome in cell culture.

### Optimized amino acid combination for pSILAC-based immunopeptidomics

While essentially all peptides from a tryptic digest can be isotopically labeled in a SILAC experiment by a combination of lysine and arginine, the percentage of HLA-I peptides that can be labeled with these two amino acids varies greatly among different HLA allotypes. In the Ma-Mel-63a immunopeptidome 54% of the HLA-A*02 and 58% of the HLA-B*18 peptides do not contain any arginine or lysine and would therefore remain unlabeled (**Supplemental Figure 3A**). In contrast, almost all peptides with basic anker residues, such as HLA-A*03-restricted peptides (96%) do contain arginine or lysine. Analyzing the amino acid composition of HLA-I peptides identified from of 96 monoallelic cell lines (26) further confirmed that the percentage of peptides that can be labeled by a combination of arginine and lysine varies widely among different HLA allotypes. On average, 28% of all peptide sequences remained unlabeled by using arginine and lysine only. For most HLA allotypes, a percentage of 20 - 70% of the HLA peptides remain unlabeled (**Supplemental Figure 4A**). We then considered using a third (conditionally) essential amino acid to increase the proportion of HLA peptides that can be metabolically labeled. An analysis of the amino acid abundances of ∼770k HLA-I peptide sequences from the IEDB database revealed that leucine is the most prevalent amino acid with 68% of the peptides containing at least one leucine, followed by valine with 51% and alanine with 50%, glutamic acid with 48% and isoleucine with 42% (**Supplemental Figure 5A**). A comparison of the peptide fractions that remained unlabeled using different amino acid combinations showed that labeling with lysine, arginine and leucine was predicted to be the most efficient. With this combination, an average of only 8% of all peptides from the IEDB remained unlabeled, whereas when using isoleucine or glutamic acid instead of leucine, a fraction of 17% and 16%, respectively, remained unlabeled (**Supplemental Figure 5B-D**). From these analyses, we concluded that lysine, arginine and leucine is the best amino acid combination for pSILAC-based immunopeptidomics experiments, resulting in an average percentage of 92% isotopically labeled HLA-I over all HLA allotypes (**Supplemental Figure 4B/ Supplemental Figure 5D**).

We decided to use isotopically labeled leucine-3 (D_3_-Leu; Leu3) for the medium-heavy state (M) and leucine-7 (^13^C_6_,^15^N_1_-Leu; Leu7) for the heavy state (H). However, in initial experiments with these isotopomeres we observed a partial loss of the ^15^N label from the α-amino group of Leu7 in leucine-containing peptides (**Supplemental Figure 6**). While Leu7 partially lost its ^15^N label, ^15^N-labeled arginine (Arg10) and ^15^N-labeled lysine (Lys8) maintained their ^15^N label. We assumed that the loss of the ^15^N label specifically from Leu7 is caused by a transamination reaction catalyzed by the enzyme branched-chain amino acid transaminase 1 (BCAT1), which seems to be more active than transaminases that act on Lys and Arg. Of note, BCAT1 is often overexpressed in leukemia cells (27) and potentially also in other tumor cells, making the use of Leu7 for quantitative immunopeptidomics even more problematic. We therefore decided to use Leu6 instead, which still shows a 3 Da mass difference to Leu3 but does not harbor a 15N label at its α-amino group.

### Optimized data analysis strategy improves accuracy of quantitative immunopeptidomics

To assess the quantification accuracy of the established pSILAC-based workflow, we performed control experiments (mock) with 0.1% DMSO treatment during the heavy and the medium-heavy labeling pulse for 24h and for 72h. We analyzed the data with Peptide-PRISM and MaxQuant as described above and observed that H/M ratios from MaxQuant evidence table of type ISO-MSMS (i.e. peptide features for which only one labeling state has been detected) showed increased ratio scattering compared to all other types (not ISO-MSMS) (Figure 2A). Especially when the medium-heavy (=1) or heavy (=2) labeling state is missing (Figure 2A; type ISO-MSMS, open circles), quantification becomes less accurate, since in these cases, MaxQuant calculates the ratios from peak intensity vs. background (re-quantify option). In addition, we realized that some peptide features have not been recognized by MaxQuant as isotope clusters and are classified as type ISO-MSMS, although both labeling states have been detected. We noticed that quantification of these peptide features becomes more error-prone, when the peptide sequences contain three or more Lys and/or Leu residues (Figure 2A, type ISO-MSMS, red filled circles), as for example for the peptide GLLGTLVQL. This peptide contains four Leu residues und thus 12 deuterium atoms. According to the reduced hydrophobicity of deuterium compared to hydrogen (27), the medium-heavy-labeled isotopomere of GLLGTLVQL showed a large retention time shift of -0.3 minutes (Supplemental Figure 7), which seems to impair the recognition of isotope clusters.

We found that calculating H/M ratios from the heavy and medium intensities given in the MaxQuant evidence table results in improved quantification accuracy for peptide features of type ISO-MSMS when both labeling states (0,1,2 or 1,2) have been detected (Figure 2B).

Finally, we calculated the “Number of reliable and corrected ratios”, from which the median peptide ratio was calculated as a measure of the expected quantification accuracy (see Methods for details). This parameter allows to recognize peptides, for which the H/M peptide ratio calculation is based solely on unreliable H/M ratios. These were typically detected with low intensities and show increased ratio scattering (Figure 2D, white circles). Both, the correction of H/M ratios for peptides with many Leu and/or Lys residues as well as the introduction of parameters for assessing quantification accuracy improved the outcome of the quantitative immunopeptidome analysis. (Figure 2C **vs. 2D**). This is further exemplified by the PRAME-derived HLA peptide SLLQHLIGL, which appears to be falsely upregulated in the control experiment when using the original H/M ratios from MaxQuant peptides table (Figure 2C), but unregulated when using the corrected H/M ratios calculated from MaxQuant evidence table (Figure 2D).

For all subsequent treatment experiments, we performed a control experiment for each cell line as described above and used the interquartile-range (IQR) to define significance levels (1.5-fold IQR and 3-fold IQR) to identify HLA peptides modulated by the corresponding treatment (**Supplemental Figure 8**).

### Short-term inhibition of MAPK pathway modulates the immunopeptidome

Next, we applied the optimized pSILAC workflow to study how the immunopeptidomes of two BRAF^V600^-mutated melanoma cell lines, Ma-Mel-63a and UKE-Mel-105b, are modulated by pharmacological inhibition of the MAPK pathway. For this purpose, we employed the (i) MEK inhibitor trametinib and (ii) vemurafenib plus cobimetinib, an FDA- and EMA-approved BRAF/MEK inhibitor combination for BRAF^V600^-mutated melanoma.

Concentrations of 100 nM trametinib and 1 µM vemurafenib plus 100 nM cobimetinib led to profound inhibition of the MAPK pathway (**Supplemental Figure 9**) and thus were applied for all quantitative immunopeptidomics experiments described below. On average we identified ∼5000 conventional HLA-I peptides derived from coding sequence (CDS) and ∼500 cryptic HLA-I peptides for Ma-Mel-63a and for UKE-Mel-105b with Peptide-PRISM (category- and length-specific FDR < 10%). The high percentage (∼10%) of cryptic peptides is due to the expression of HLA-A*03:01 in both cell lines. The immunopeptidomes of both melanoma cell lines exhibited substantial quantitative changes upon treatment with trametinib. ∼10% of all HLA-I peptides showed significantly increased and another ∼10% significantly decreased presentation, based on the 3-fold interquartile range (IQR) of the corresponding mock experiment (**Supplemental Figure 10**). We found that peptides from melanoma differentiation antigens (MDAs, **Supplemental Table 4**), such as GTYEGLLRR and SLDDYNHLV (dopachrome tautomerase, DCT), ALDGGNKHFL, ALLAVGATK (premelanosome protein, PMEL) and ALLAGLVSL (tyrosinase, TYR) were consistently upregulated by trametinib (Figure 3E-H), which is in line with the previously described upregulation of MDAs early under MAPKi treatment in Me-Mel-63a (28) and which is further in accordance with results published using a TMT-labeling approach (6).

Next, we aimed for identifying further tumor-associated or -specific T cell epitopes that are linked to an overactivated MAPK pathway. To classify HLA peptides as tumor-exclusive, we re-analyzed a large set of publicly available immunopeptidome data from tumor and healthy samples (for details see Methods) and identified conventional and cryptic HLA peptides with Peptide-PRISM. In addition, we analyzed the immunopeptidome of commercially available, primary melanocytes. Since melanoma originates exclusively from melanocytes or their progenitor cells, melanocytes should be the best reference to identify truly melanoma-specific HLA peptides. To maximize the overlap of HLA peptides with the two melanoma cell lines, we chose HLA-A*03:01-positive melanocytes. From the combined data we generated a database (tumor vs. healthy database, TvHdb) by mapping all identified HLA peptides to their corresponding open reading frames (ORFs) and classified those ORFs as tumor-exclusive, from which HLA peptides were detected only in tumor samples. With the information from TvHdb, we classified all HLA peptides from pSILAC experiments that derived from tumor-exclusive ORFs as “tumor” and all peptides from non-tumor-exclusive ORFs as “tumor & benign” (Figure 3). In addition, we extracted from the TvHdb in how many different patients or cell lines the corresponding peptides have been detected and from which sample types (tumor entity or origin of healthy sample; **Supplemental Tables 5**).

As expected, the great majority of HLA peptides were not tumor-exclusive and probably represents self-peptides. HLA peptides from MDAs, such as DCT, PMEL and tyrosinase were recurrently detected exclusively in melanoma and melanocytes. One putative tumor-specific peptide with the sequence TTNARILAR, derived from urocortin-2 (*UCN2*) was downregulated upon MAPK-signaling inhibition in Ma-Mel-63a cells. This protein is known to activate ERK1/2 signaling and plays a role in the context of cardio protective effects (29). It was identified in 6 out of 12 HLA-A*03:01 positive melanoma patients. Transcriptomic analyses did not reveal any evidence for tumor-specific expression pattern of this gene (30). We therefore decided to not assess immunogenicity of this conventional HLA peptide. Essentially, no other tumor-exclusive HLA peptides were repeatedly detected in the pSILAC data from MAPK inhibition experiments. However, we reproducibly identified two tumor-exclusive cryptic peptides that changed upon MAPK pathway inhibition. The peptide RVYLDIVTPK (off-frame, HLA-A*03:01, *IFT81*) was reproducibly upregulated under all tested conditions and the peptide RLSSSLPSR (5’UTR, HLA-A*03:01, *ST3GAL6*) was reproducibly downregulated upon MAPK pathway inhibition in Ma-Mel-63a-cells (Figure 3 and **Supplemental Figure 11** and **Supplemental Figure 12**). In TvHdb RVYLDIVTPK (*IFT81*) was identified exclusively in melanoma samples (in 17 of 31 HLA-A*03:01/A*11:01/A*31:01-positive tumor metastases or patient-derived cell lines) and RLSSSLPSR (*ST3GAL6*) was identified in melanoma (in 9 of 31 HLA-A*03:01/A*11:01/A*31:01-positive melanoma samples) (**Supplemental Table S6**). This could indicate a relevance of these peptides particularly in melanoma disease and progression. RVYLDIVTPK is an off-frame peptide, deriving from the coding region of *IFT81* in a non-canonical reading frame (**Supplemental Figure 13**). Ribosome profiling data (nuORF database (31)) indicated that this peptide derives from a short (34 amino acids) internal ORF with an alternative CTG start codon. The protein IFT81 (protein intraflagellar transport 81) is involved in cilia assembly (31). Interestingly, Jenks *et al.* described changes in cilia length and formation as a hallmark of (MEK) kinase inhibitor resistant cells (32). This supports our hypothesis, that the generation of the cryptic peptide RVYLDIVTPK (*IFT81*) could be linked to inhibited MAPK signaling and could derive from activated alternative signaling pathways. Although the exact mechanism still needs to be unraveled, our studies strongly indicate that the cryptic peptide RVYLDIVTPK is generated upon MEK-inhibitor exposure. According to the nuORF database, the cryptic peptide RLSSSLPSR (*ST3GAL6*) derives from an upstream open reading frame (uORF) also with an alternative CTG start codon (**Supplemental Figure 14**). uORFs can regulate expression of downstream gene products. The translation of the corresponding uORF, associated with the generation of RLSSSLPSR could therefore suppress translation of ST3GAL6. A tumor suppressive role of ST3GAL6 (Type 2 lactosamine alpha-2,3-sialyltransferase) was indeed described by Li *et al*. (33), which could indicate that suppressed ST3GAL6 levels could be advantageous for tumor progression. Targeting this peptide under non-treatment conditions could be a promising immunotherapeutic strategy, provided that the tumor relies on ST3GAL6 suppression and the generation of the RLSSSLPSR peptide as byproduct of this tumor promoting mechanism.

### Testing immunogenicity of cryptic peptides modulated by MAPK pathway inhibition

To collect further evidence for the tumor specificity of the two cryptic peptides RVYLDIVTPK (*IFT81*) and RLSSSLPSR (*ST3GAL6*), we evaluated their immunogenicity applying an *in vitro* priming assay (Figure 4). In addition to the two HLA-A*03:01-restircted cryptic peptides, we included one HLA-A*02:01- and one A*03:01-restricted peptide from DCT that showed increased presentation upon MAPK pathway inhibition. For *in vitro* priming we used PBMCs of two independent HLA-A*02- and HLA-A*03-positive healthy donors. The two self-peptides TLYEAVREV (HLA-A*02) and RILGPGLNK (HLA-A*03) derived from RPL10A with expected central tolerance were used as negative control. The peptides ELAGIGILTV (MART-1, HLA-A*02), YLQPRTFLL (SARS-CoV2, HLA-A*02) and ILRGSVAHK (Flu, HLA-A*03) served as positive controls.

**Figure 4.**
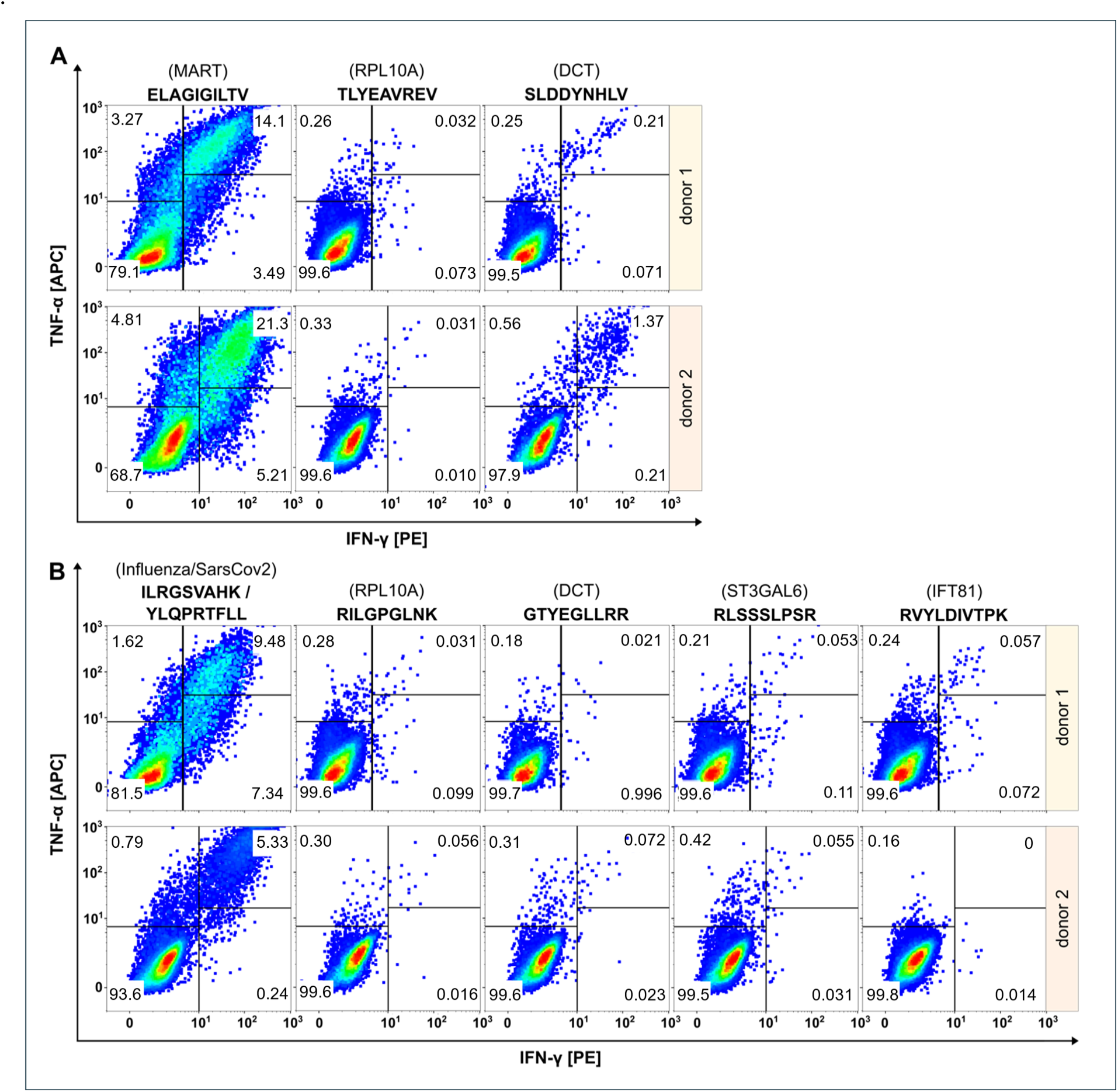
Evaluation of immunogenicity of HLA peptides regulated upon BRAF and/or MEK inhibition by *in vitro* priming using human PBMCs from two healthy donors. Peptides restricted to HLA-A*02 are shown in (**A**), peptides restricted to HLA-A*03 in (**B**). Peptides derived from RPL10A were used as negative controls. Peptides from MART-1, SARS-CoV2 or Flu served as positive controls. The peptide SLDDYNHLV (*DCT*, HLA-A*02) induced immunological responses in both donors, while the peptide GTYEGLLRR (*DCT*, HLA-A*03) induced no immunological responses. Cryptic peptides RLSSSLPSR (*ST3GAL6*) and RVYLDIVTPK (*IFT81*) elicited only marginal immunological response.

Effector responses were evaluated by gating TNFα/IFNγ double positive cell populations of CD8^+^ T cells. The frequency of TNFα/IFNγ-positive T cells in all negative controls was below 0.06% and ranged between 5.33-21.3% for the positive controls. The HLA-A*02:01-restricted peptide SLDDYNHLV (derived from DCT) induced immunological responses in both healthy donors with fractions of 0.22% and 1.37% double positive cells, respectively, confirming the immunogenicity of this peptide as described earlier (34). In contrast, the HLA-A*03-restricted peptide GTYEGLLRR that also derived from DCT, induced fractions of only 0.021% and 0.072% double positive cells that were almost equal to negative control level. Cryptic peptides RLSSSLPSR (ST3GAL6) and RVYLDIVTPK (IFT81) showed only marginal responses in donor 1 (0.053% and 0.057% versus 0.031% in the negative control) and no responses in donor 2 (0.055% and 0% versus 0.056% in the negative control).

### Resistance against MAPKi results in substantial immunopeptidome alterations

Long-term therapeutic efficacy of MAPKi is often limited due to acquired drug resistance and a major cause for clinical tumor progression or relapse. There are controversial results from randomized phase III studies concerning the anti-tumor activity of the combined treatment of BRAF-mutated metastatic melanoma with immune checkpoint and MEK inhibitors (35). In this context, we aimed to study how drug resistance contributes to immunopeptidomic alterations that can impact tumor immunogenicity. To this end we generated a MEKi-resistant melanoma cell line. Since Ma-Mel-63a cells that were long-term exposed to BRAFi and MEKi (Ma-Mel-63a dr) turned out to be still sensitive to BRAF and MEK inhibition (**Supplemental Figure 15**), we induced BRAFi- and MEKi resistance in the BRAF^V600^-mutated melanoma cell line SK-Mel-28 by long-term exposition to vemurafenib and cobimetinib. In contrast to SK-Mel-28 wt, the MAPK pathway was no longer inhibited by treatment with vemurafenib/cobimetinib for SK-Mel-28 dr (Figure 5A).

**Figure 5.**
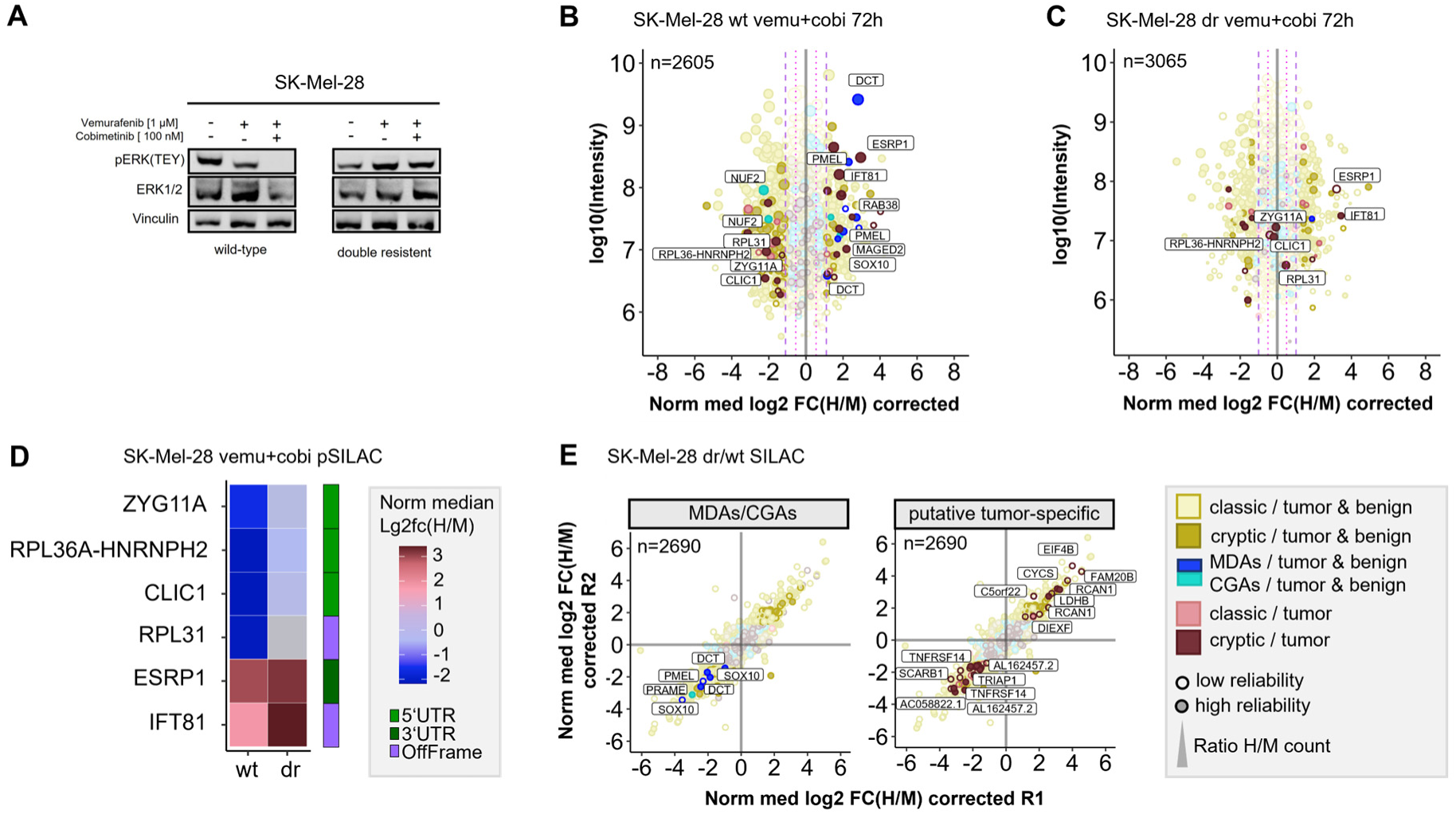
Quantitative Immunopeptidome analyses of SK-Mel-28 wt and SK-Mel-28 dr cells. (**A**) Western blot analysis confirmed inhibited MAPK signaling in SK-Mel-28 wt cells and active MAPK signaling in SK-Mel-28 dr cells upon BRAF- and MEK-inhibitor treatment. (**B**) Pulsed-SILAC based immunopeptidome analysis of SK-Mel-28 wt cells upon treatment with vemurafenib and cobimetinib. HLA presentation is globally altered, and MDAs show enhanced presentation. (**C**) Immunopeptidomes of SK-Mel-28 dr cells also showed alterations, however, MDA presentation was not altered and only few cryptic peptides reproducibly changed their presentation upon kinase inhibitor treatment. Mock control experiments of SK-Mel-28 dr were used to determine significantly regulated peptides. (**D**) Modulation of putative tumor-specific cryptic peptides under basal conditions and upon kinase inhibitor treatment in SK-Mel-28 wt and SK-Mel-28 dr cells. Peptides, that were downregulated in wild-type cells with inhibited MAPK signaling but not in SK-Mel-28 dr cells with maintained MAPK signaling could directly be attributed to active MAPK signaling pathway. Peptides that were upregulated in both cell lines could be directly linked to the kinase inhibitors but not to MAPK signaling. (**E**) Immunopeptidome changes of resistant and wild-type SK-Mel-28 cells. Presentation of MDAs (*PMEL, DCT, SOX10*) is reduced in SK-Mel-28 dr cells. Presentation of some putative tumor-specific cryptic peptides is rearranged.

First, we conducted pSILAC experiments with SK-Mel-28 wt and SK-Mel-28 dr cells to unravel short-term immunopeptidomic changes upon BRAF- and MEK inhibition with a combination of vemurafenib and cobimetinib over 24h and 72h. In analogy to melanoma cell lines Ma-Mel-63a and UKE-Mel-105b, SK-Mel-28 wt cells showed enhanced presentation of MDA-derived peptides upon MAPK pathway inhibition (Figure 5B). In contrast, SK-Mel-28 dr cells did not show this effect (Figure 5C). In addition, we recognized that two HLA peptides from NUF2 were downregulated upon MAPK pathway inhibition in SK-Mel-28 wt cells (Figure 5B). NUF2 exhibits a Cancer germline-like expression profile in GTEx (36) with predominant expression in testis and marginal expression in most other tissues. Expression of Cancer germline antigens (CGAs) get lost when stem cells differentiate and can be reactivated during oncogenesis when tumor cells dedifferentiate.

Next, we focused on cryptic peptides. We observed that many of these showed an altered presentation in vemurafinib/cobimetinib-treated SK-Mel-28 wt cells (Figure 5B). In contrast, in SK-Mel-28 dr cells, only a few cryptic peptides showed an altered presentation (Figure 5C) and most of these were not reproducibly detected for both time points (**Supplemental Figure 18**). We identified two (according to TvHdb) putative tumor-specific cryptic peptides (RVYLDIVTPK, *IFT81* and SSIRSWNNK, *ESRP1*) with increased presentation after short-term MAPKi treatment in wildtype as well as in resistant cells (Figure 5B **& C**), indicating that their presentation is not directly linked to MAPK signaling. Moreover, we identified four putative tumor-specific cryptic peptides from *ZYG11A, RPL36A-HNRNPH2, CLIC1* and *RPL31* that were decreasingly presented upon MAPKi in SK-Mel-28 wt but were unaffected in SK-Mel-28 dr cells (Figure 5B **& C**), indicating a direct link to MAPK signaling. A linkage to MAPK signaling has already been reported for one of these genes, chloride intracellular channel 1 (*CLIC1*), in the context of gastric cancer progression (37). All four HLA peptides have been recurrently identified in different tumor entities but never on healthy cells in TvHdb (**Supplemental Table 6**). For instance, the peptide SVASTNPIK (*RPL36A-HNRNPH2)* has been identified in 10 tumor patients (melanoma, CRC, GBM), representing a recurrent putative tumor-specific T-cell antigen.

Subsequently, to study long-term alterations in the immunopeptidome that are linked to MAPK inhibitor resistance, we conducted a SILAC-based experiment to directly compare the immunopeptidomes of SK-Mel-28 wt and SK-Mel-28 dr cells. We found that formation of resistance was accompanied by substantial immunopeptidomic alterations, reflecting changes of the cellular state. We observed that the overall presentation of HLA peptides derived from MDAs (*SOX10, PMEL, DCT*) was diminished in SK-Mel-28 dr cells compared to SK-Mel-28 wt cells (Figure 5E), indicating an epithelial-to-mesenchymal-like transition during long term MAPKi exposure. We found additional evidence for this phenotype switch of drug-resistant cells towards a dedifferentiated cell state by RT-qPCR (**Supplemental Figure 19A**) and by proteome analysis (**Supplemental Figure 19B**). On transcriptome level, SK-Mel-28 dr cells showed elevated AXL and reduced MITF expression, representing a trend towards an AXL^high/^MITF^low^ phenotype, which defines a dedifferentiated melanoma phenotype (38). While differentiated melanomas highly express MDAs, expression of MDAs is lost in dedifferentiated melanomas (39). Accordingly, we observed decreased protein levels of MDAs (*DCT, PMEL*) in the proteome of SK-Mel-28 dr compared to SK-Mel-28 wt cells. The observed diminished presentation of HLA peptides from MDAs suggests that patients with acquired MAPK inhibitor resistance might show decreased responsiveness to immunotherapeutic treatments targeting MDAs.

After that, we found that a number of putative tumor-specific cryptic peptides showed significantly increased presentation in MAPKi-resistant cells, while others exhibited substantially decreased presentation (Figure 5E). The peptides with increased presentation in SK-Mel-28 dr cells derived from *EIF4B, FAM20B, LDHB, CYCS, C5orf22, RCAN1* and *DIEXF* and were recurrently identified in TvHdb in a variety of tumor patients **(Supplemental Table 6)**. For example, the peptides RVQILQILK (*DIEXF*) and STDLPILLK (*C5orf22*) were identified in 9 and 7 tumor patients, respectively. In general, upregulation of these cryptic peptides might mechanistically be explained by adaptive response of the tumor cells to stress conditions, such as pro-apoptotic or hypoxic conditions, induced by kinase inhibitor exposition. As a result, the expression of genes that adapt the cell state to overcome these unfavorable cellular conditions, is altered and the transcriptional landscape is changed. For instance, translation of off-frame cryptic peptides from transcripts of lactate dehydrogenase *(LDHB*) and cytochrome c (*CYCS*) could be a result of ongoing metabolic alterations to overcome kinase inhibitor-induced cell stress. Contretras Mostazo *et al.* reported that *CYCS* was upregulated in an imatinib-resistant CML cell line, confirming that mitochondrial respiration can be influenced by kinase inhibitor exposition (40). *RCAN1* contributes to deregulation of calcineurin-involved signaling pathways, however, its role to cancer progression is contradictory. In melanoma, transcript levels of *RCAN1* are elevated in comparison to normal samples, indicating that there might be an association to melanoma progression (41). Overall, putative tumor-specific cryptic peptides with enhanced presentation on kinase inhibitor-resistant melanoma cells, might represent attractive targets for immunotherapy, especially in combination with MAPKi treatment.

In contrast, putative tumor-specific cryptic peptides with diminished presentation on MAPKi-resistant melanoma cells are probably less attractive targets, but still reflect alterations of the cellular state. We identified cryptic peptides with reduced presentation on MAPKi-resistant cells from *TRIAP1, SCARB1, AC058822.1, AL162457.2* and *TNFRSF14* (Figure 5E, **Supplemental Table 6**). Two of these peptides derived from proteins that are related to necroptosis. Necroptosis is an alternative type of programmed cell death, in which processes of necrosis are involved. Necroptosis-related cryptic peptides derived from *TNFRSF14*, a TNF receptor gene and a long-non-coding RNA (lncRNA) AL162457.2. Reports of Liu *et al.* (42) indicate decreased overall survival in melanoma when this necroptosis-related lncRNA is expressed. Interestingly, findings of Sugaya *et al.* (43) show that the BRAF inhibitor dabrafenib attenuates necroptosis. These results combined with our data suggest, that necroptosis and expression of necroptosis-related genes is inhibited during long-term BRAFi exposure, finally leading to a decreased presentation of the corresponding cryptic HLA peptides.

## Discussion

In this work, we have successfully established a pSILAC-based approach for quantitative immunopeptidomics. We have demonstrated that our strategy, combining *de novo* sequencing-based HLA peptide identification with Peptide-PRISM and quantification with MaxQuant, enables to monitor modulations of the immunopeptidome with high accuracy. Moreover, this approach enables for the first time to quantify variations in the presentation of cryptic HLA peptides. This opens up the possibility to improve our understanding of this so far insufficiently characterized class of HLA peptides. The workflow is simple to conduct and ideally suited to study drug-induced alterations of the immunopeptidome in cell culture. We have demonstrated this by analyzing how different MAPK signaling pathway inhibitors change the immunopeptidome of various melanoma cells lines. This method can be easily applied to study the effects of any drug, such as epigenetic or cytostatic drugs, on the immunopeptidome. In addition, this approach can also readily be used to study the effect of specific protein knockdowns (e.g. by inducible shRNA expression), protein degradations, accomplishable for example by the PROTAC approach (44) or ectopic expression of specific proteins, such as viral immunoevasins (45). Furthermore, we have demonstrated that a SILAC-based approach, using complete instead of pulsed labeling, enables an accurate quantitative comparison of the immunopeptidomes of isogenic cell lines. Together, this allowed us to uncover profound differences in the immunopeptidomes induced upon short-term MAPKi-treated cells and upon acquired MAPKi resistance.

So far, immunopeptidomes were often compared in a qualitative way by comparing lists of identified HLA peptides. Considering the typically moderate immunopeptidome coverage, the limited reproducibility of the affinity purification, and associated with both, the large number of missing values, this is a quite error-prone procedure. One major advantage of SILAC-based immunopeptidomics compared to label-free or isobaric labeling is that the two samples to be compared can be pooled already before cell lysis. This excludes inaccuracies introduced by separately performed affinity purifications and substantially improves quantification accuracy. Quantification accuracy is of utmost importance for quantitative immunopeptidomics, since peptide quantification is much less error-tolerant than protein quantification. Data-independent analysis (DIA) of SILAC-based immunopeptidomics samples might enable even higher quantification accuracy (46). On the downside, SILAC-based immunopeptidomics is basically limited to a maximum of three samples that can be directly compared in one analysis.

Using SILAC-based quantitative immunopeptidomics allowed us to identify various cryptic HLA peptides, the presentation of which is altered early in response to MAPKi treatment or on MAPKi-resistant cells. Especially tumor-exclusive cryptic peptides with enhanced presentation on MAPKi-resistant cells might represent attractive targets for cancer immunotherapy. These examples demonstrate that alterations of the cellular state are reflected by cryptic HLA peptides at least equally well as by conventional HLA peptides from CDS. Although we could not detect substantial T cell reactivities in healthy donor PBMCs toward the two cryptic HLA peptides in our experimental *in vitro* priming setting, this does not exclude their immunogenicity in cancer patients. If it is not possible to isolate high affinity TCRs against these peptides by *in vitro* priming with healthy donor cells, generating high affinity TCRs in mice (47) or generating peptide-centric CARs (48) might be alternative approaches.

We envision a large range of applications for SILAC-based quantitative immunopeptidomics. For instance, this method can be applied to study the effects of oncogene overexpression or the expression of proteins with oncogenic driver mutations, such as histone H3K27M in diffuse midline glioma. This might reveal T-cell epitopes whose presentation is directly linked to driving oncogenic activity. Such T-cell epitopes would presumably represent promising targets for cancer immunotherapy. By using isotopically labeled cell lines, or mixtures of cell lines, as for the Super-SILAC approach (49), SILAC-based quantitative immunopeptidomics might also be valuable for detecting differences between immunopeptidomes of healthy and tumor tissue. To the same extent as for tumor immunity, we can imagine various applications in the field of autoimmunity and antiviral immunity. Having this large variety of possible applications in mind, we expect that SILAC-based quantitative immunopeptidomics will become a widely used approach.

## Data Availability

The mass spectrometry proteomics data have been deposited to the ProteomeXchange Consortium via the PRIDE (50) partner repository with the dataset identifier PXD054097.

## Acknowledgements

This work was supported by Deutsche Forschungsgemeinschaft (DFG) research grants SCHL 1888/7-1, SCHL 1888/8-2, SCHI 1344/4-1, ER 927/6-1. Parts of this project and Anne Rech were supported by the Novartis Melanoma Award for Young Academics (MAYA 2021).

## Author Contributions

M. Bernhardt and A. Rech‡: performed experiments, data analysis, visualization, manuscript writing. M. Berthold: performed experiments. M. Lappe: performed experiments. JN. Herbel: performed experiments. F. Erhard: software development, reviewed and edited manuscript. A. Paschen: scientific concept, reviewed and edited manuscript. B. Schilling and A. Schlosser*: experimental design, scientific concept, data analysis, manuscript writing.

