## Supplemental Figures for "SILAC-based quantification reveals modulation of the immunopeptidome in BRAF and MEK inhibitor sensitive and resistant tumor cells"

**Affiliations:**

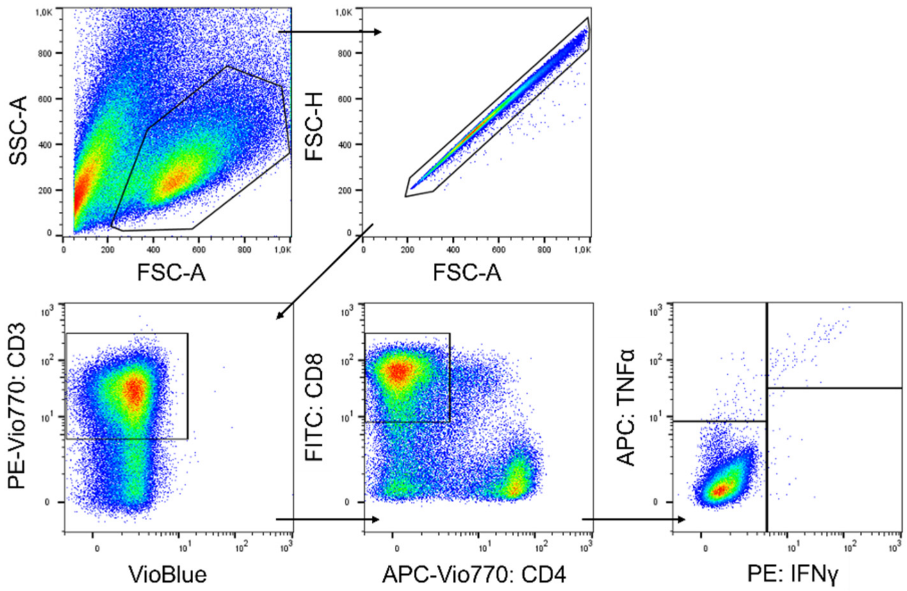


**Supplemental Figure 1 - Gating strategy for flow cytometry data after surface and cytokine staining.** $CD3^+$, $CD8^+$, $CD4^-$ T cells were selected. Data was acquired using the MACSQuant 16 Analyzer. Fluorochrome compensation was conducted following the manufacturer's instructions.

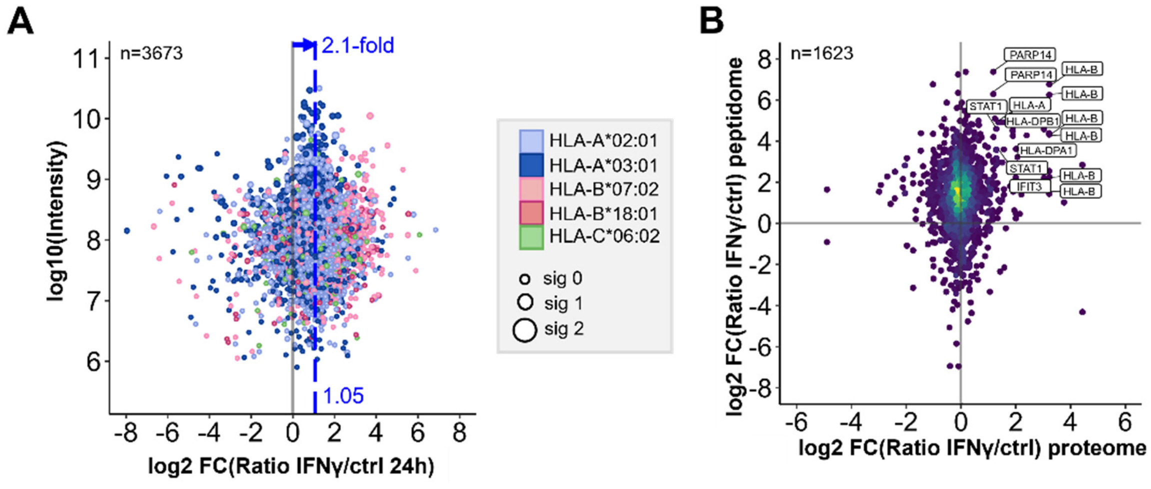


**Supplemental Figure 2 - Effects of IFNγ on the immunopeptidome and the proteome. (A)** HLA peptide presentation was globally enhanced (median 2-fold) after IFNγ for 24h. **(B)** Density correlation plot of quantitative alterations of the immunopeptidome and proteome upon IFNγ treatment. No substantial correlation was observed with exception for peptides derived from genes that were directly upregulated by IFNγ, such as genes of the antigen presenting pathway or JAK-STAT-signaling.

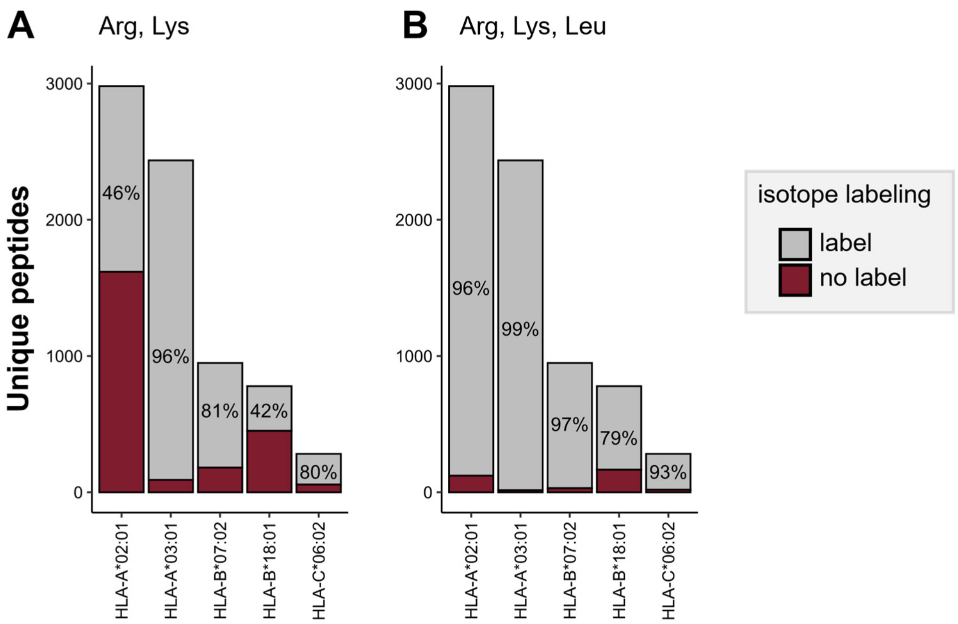


**Supplemental Figure 3 - Fractions of peptide sequences that can be isotopically labeled for the different HLA allotypes of Ma-Mel-63a.** Peptides can incorporate isotopically labeled amino acids when harboring at least one labeled amino acid per sequence. **(A)** Labeling with arginine (R) and lysine (K) results in a high percentage of labeled peptides for HLA-A*03:01, HLA-B*07:01 and HLA-C*06:01, but improper labeling for HLA-A*02:01 and HLA-B*18:01 peptides. **(B)** Labeling with R, K and leucine (L) increases the percentage of labeled peptides for all allotypes, in particular for HLA-A*02:01 and HLA-B*18:01.

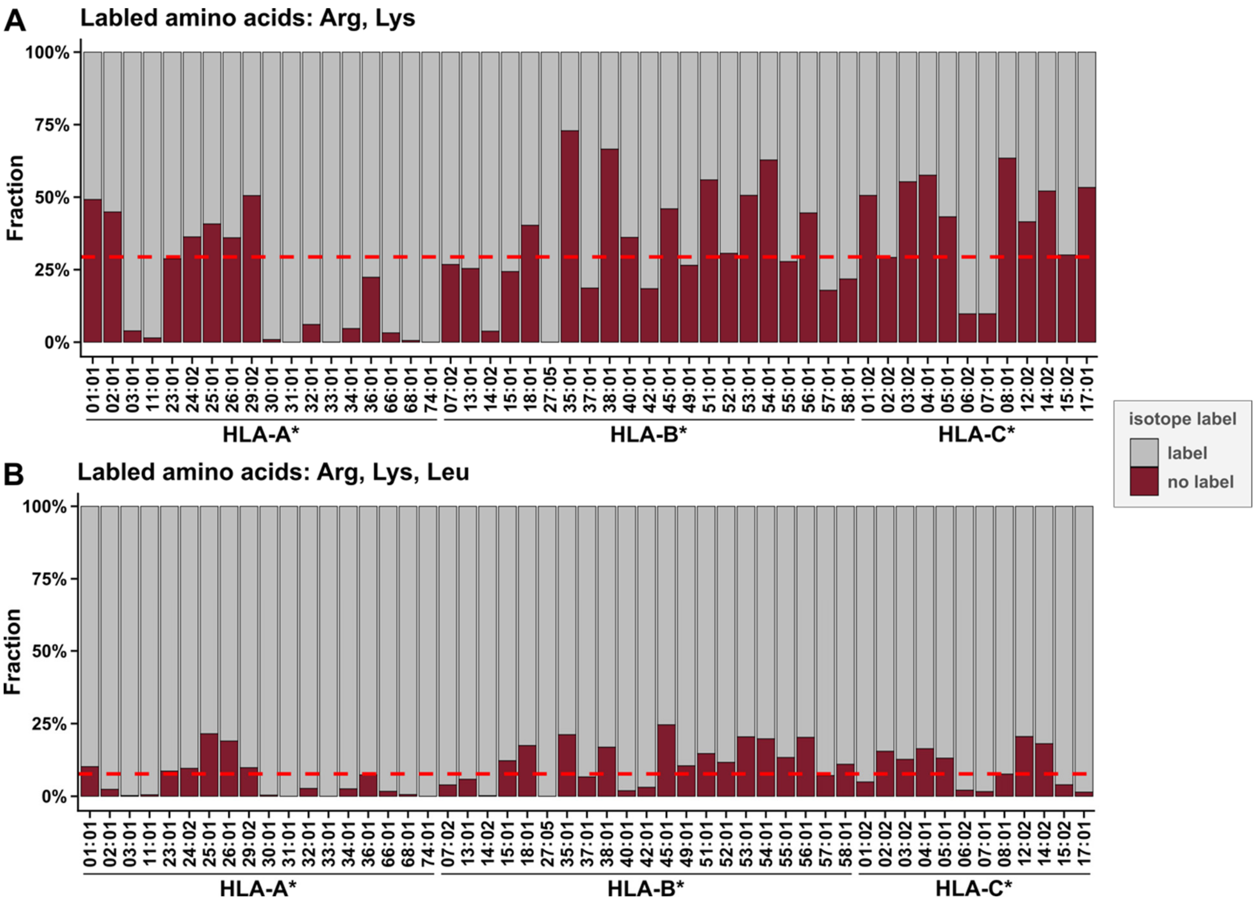


**Supplemental Figure 4 - Peptide fractions for which isotopic labeling was predicted as possible (label) or not possible (no label). (A)** The fraction of peptides that can be metabolically labeled with a combination of Arg and Lys. Insufficient labeling was predicted for most HLA-alleles. **(B)** The fraction of peptides that can be metabolically labeled with a combination of Arg, Lys and Leu. Calculations are based on a reanalysis of immunopeptidome data from 96 monoallelic cell lines with Peptide-PRISM. Identified peptides were filtered to 1% FDR and included only peptides predicted as binders by NetMHCpan 4.0.

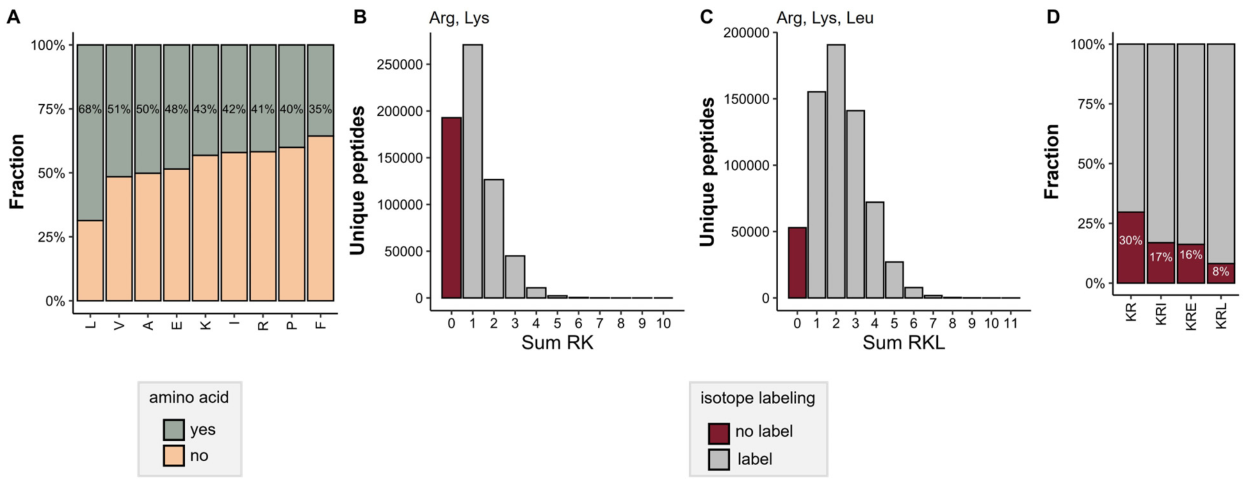


**Supplemental Figure 5 - Abundance of specific amino acids in HLA-I peptides.** Calculations are based on ~770 k HLA peptides from IEDB database. **(A)** Fraction of HLA-I peptides that contain at least one of the specified amino acids. **(B)** Distribution of peptides with no or multiple labeling amino acids in their sequence when using lysine and arginine. **(C)** Distribution of peptides with no or multiple labeling amino acids in their sequence when using lysine, arginine and leucine. **(D)** Fraction of peptides that remain unlabeled in dependence on different labeling amino acid combinations.

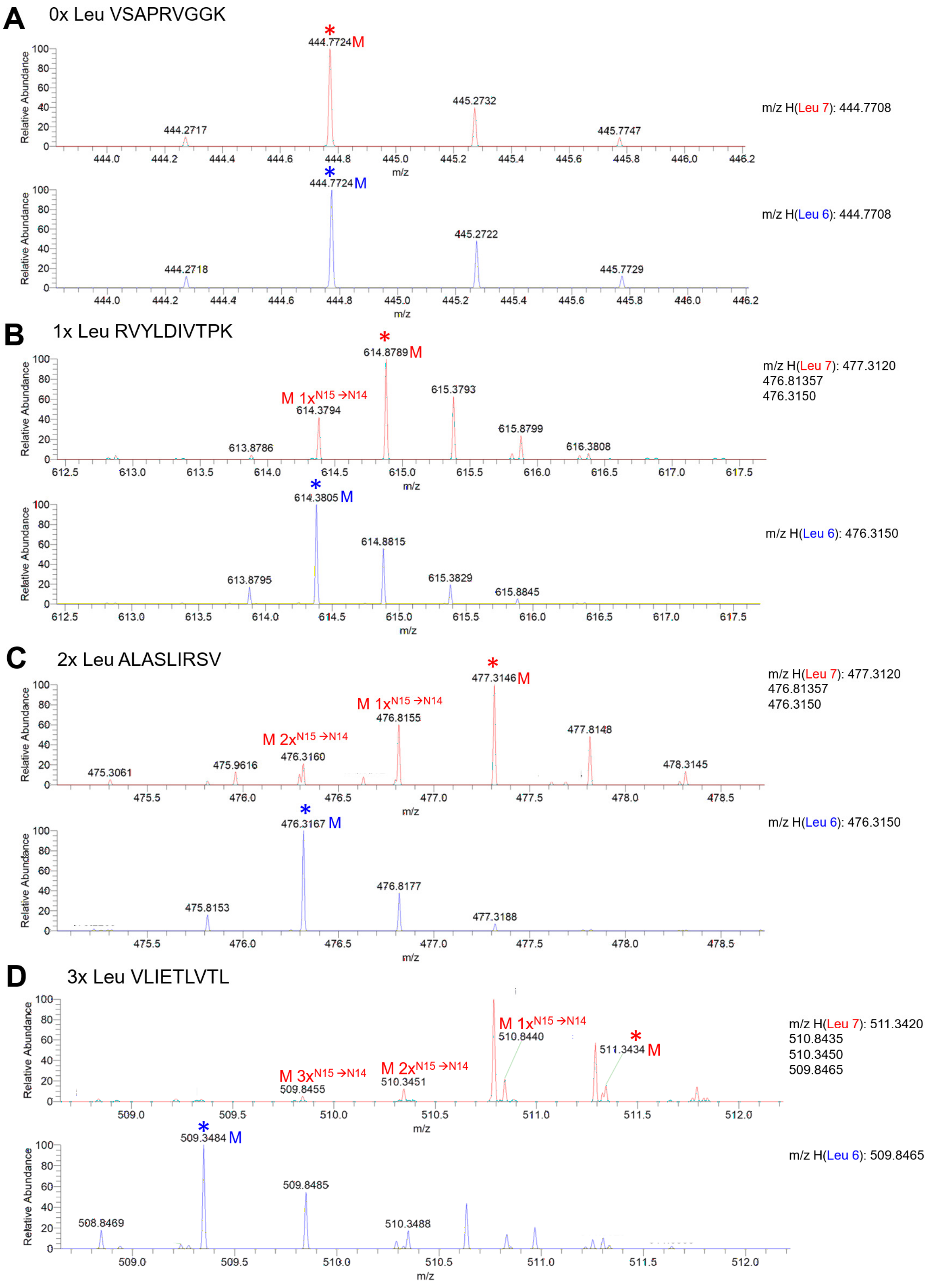


**Supplemental Figure 6 - Leucine-7 is susceptible for the loss of the 15N isotope label from the α-amino group. (A)** The peptide VSAPRVGGK without leucine residues shows identical isotopic patterns for Leu6 and Leu7 labeling. **(B)** The peptide RVYLDIVTPK with one leucine shows an additional isotopic peak at m/z =614.3794 indicative for the substitution of one 15N atom by 14N (N15 → N14) when labeled with Leu7. The same peptide labeled with Leu6 does not show any sign of isotope label loss. **(C)** The peptide ALASLIRSV with two leucine residues labeled with Leu7 generates two additional mass peaks indicative for the loss of two isotope labels (2x N15→N14). **(D)** The peptide VLIETLVTL with three leucine residues shows three additional isotopic peaks indicative the loss of three isotope labels when labeled with Leu7, but not with Leu6.

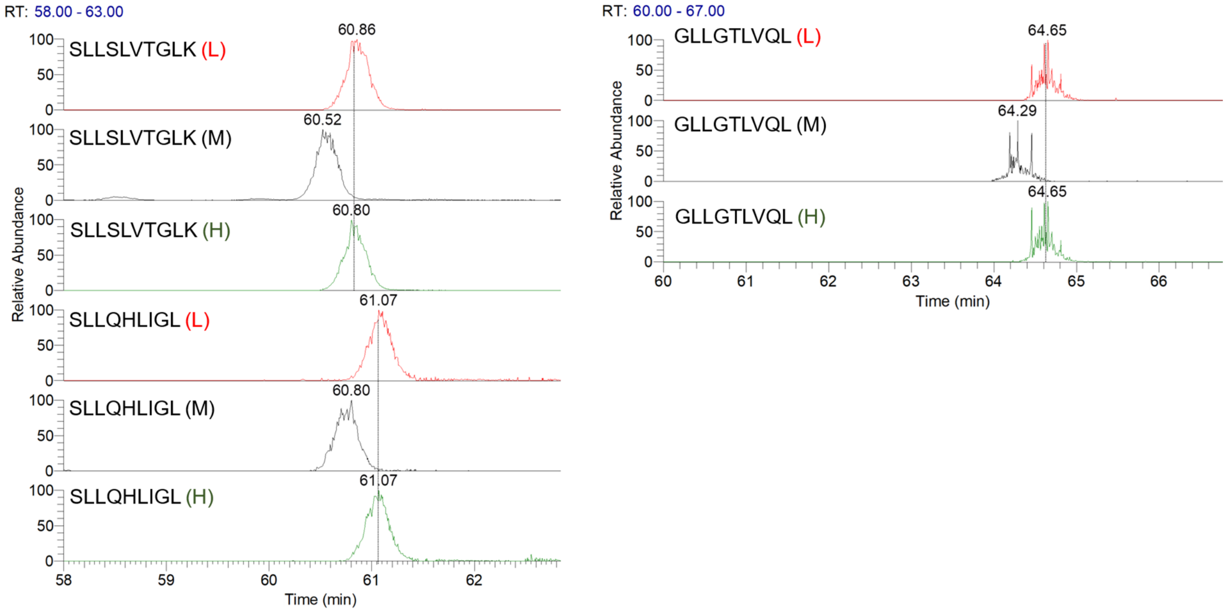


**Supplemental Figure 7 - Extracted ion chromatograms of peptides SLLSLVTGLK, SLLQHLIGL and GLLGTLVQL.** Shown peptides labeled with medium-heavy isotope amino acids elute significant earlier than light or heavy isotope-labeled counterparts, causing a retention time shift of ~ 0.2-0.3 min.

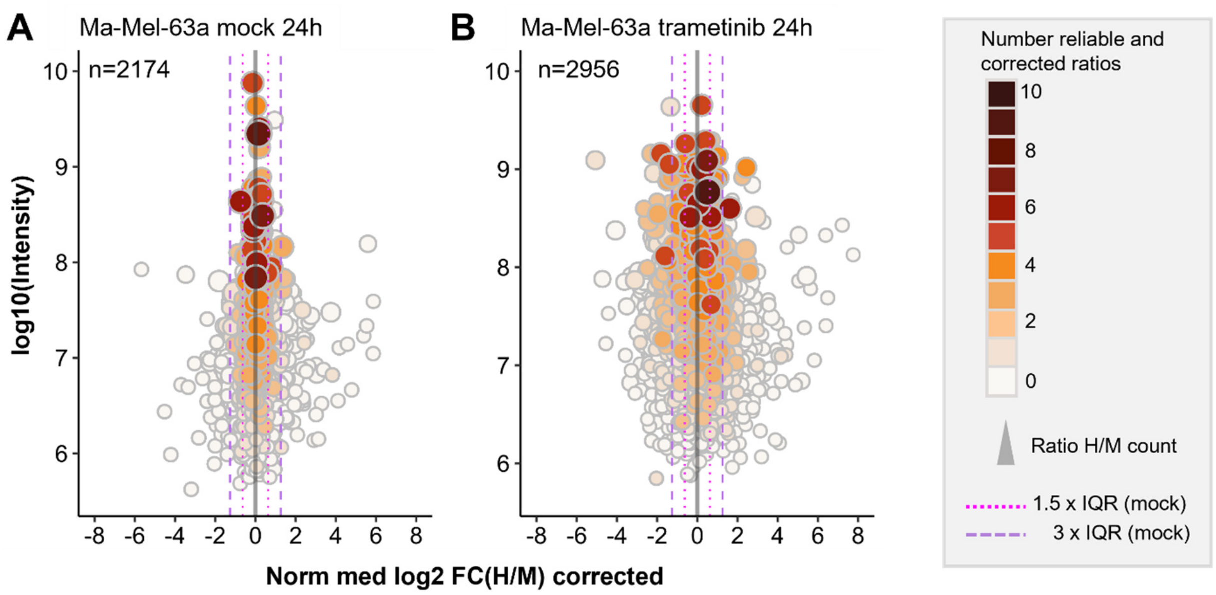


**Supplemental Figure 8 - Mock experiments were used to validate immunopeptidomic alterations occurring under treatment conditions.** Distribution of immunopeptidomic changes under mock conditions were considered to determine unspecific alterations of the immunopeptidome upon labeling. The 1.5-fold and 3-fold inter quartile range (IQR) confidence levels were used to identify background scattering. As example changes under mock conditions for Ma-Mel-63a after 24h **(A)** in comparison to changes of 24h trametinib treatment of Ma-Mel-63a **(B).**

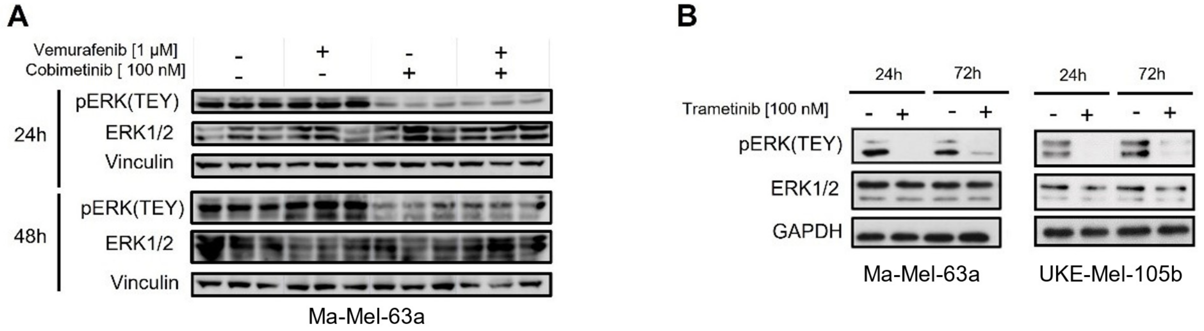


**Supplemental Figure 9 - Representative Western Blot analyses of Ma-Mel-63a and UKE-Mel-105b cells to evaluate status of MAPK signaling. (A)** Ma-Mel-63a cells were treated with vehicle (DMSO), 1 µM vemurafenib, 100 nM cobimetinib or both for either 24h or 48h. Phosphorylation of the TEY-motif of ERK was indicative for active MAPK signaling. Reduction of phospho-ERK(TEY) was observed upon MEK-inhibition (cobimetinib) alone or in combination with BRAF-inhibition (vemurafenib plus cobimetinib) but not under control conditions or BRAF inhibition after 24h and 48h. Protein levels of ERK1/2 were not altered. Vinculin was applied as loading control. **(B)** Ma-Mel-63a and UKE-Mel-105b cells were treated with vehicle (DMSO) or 100 nM trametinib for 24h and 72h. Levels of phospho-ERK(TEY) were reduced upon trametinib treatments but not under control conditions. Protein levels of ERK1/2 were not changed. GAPDH was used as loading control.

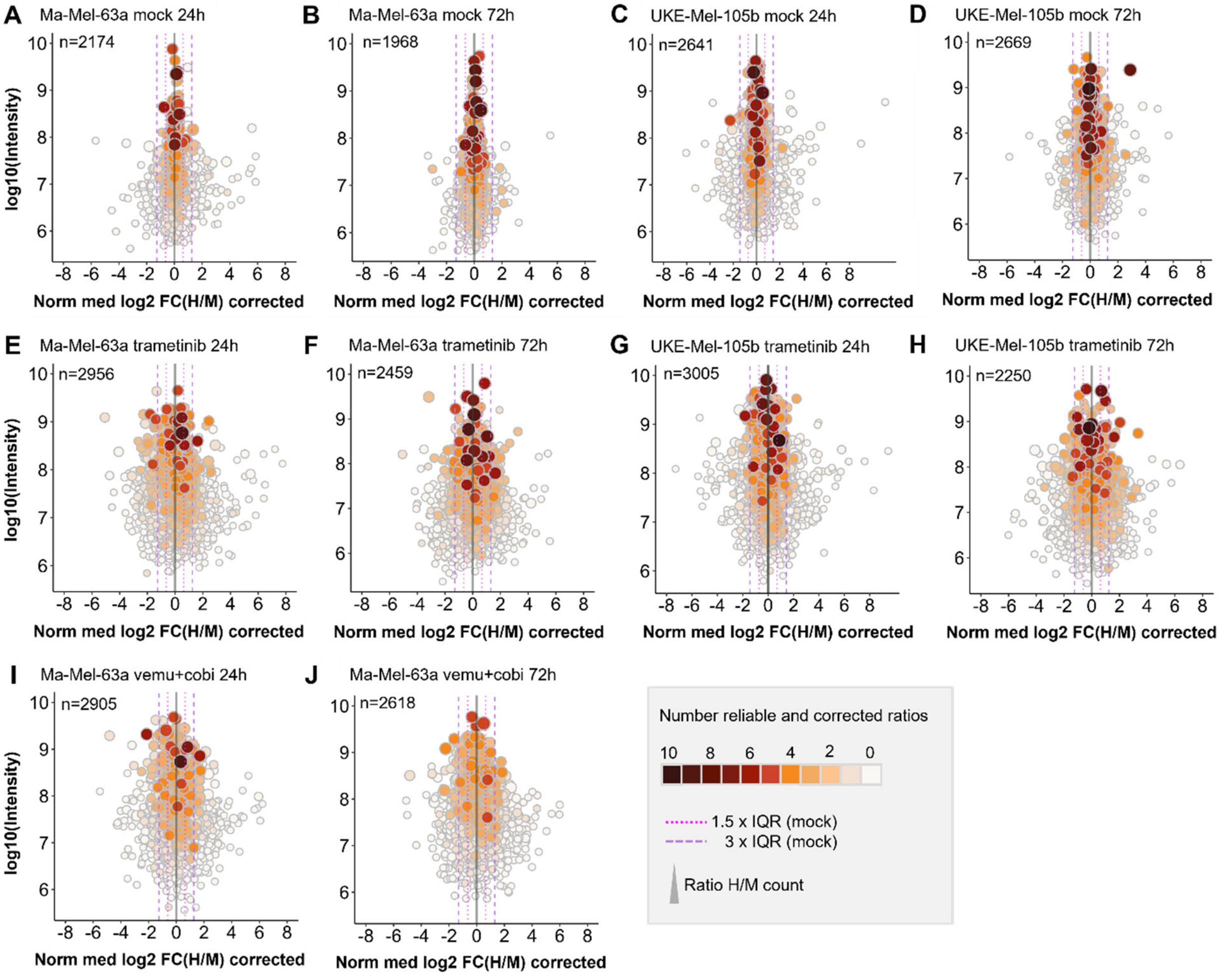


**Supplemental Figure 10 - Global immunopeptidome alterations of all tested conditions and cell lines after conducting pSILAC. (A)** Changes of the Ma-Mel-63a immunopeptidome upon 24h under mock conditions. **(B)** Changes of the Ma-Mel-63a immunopeptidome upon 72h under mock conditions. **(C)** Changes of the UKE-Mel-105b immunopeptidome upon 24h under mock conditions. **(D)** Changes of the UKE-Mel-105b immunopeptidome upon 72h under mock conditions. **(E)** Trametinib induced immunopeptidome alterations in Ma-Mel-63a cells after 24h. **(F)** Trametinib induced immunopeptidome alterations in Ma-Mel-63a cells after 72h. **(G)** Trametinib induced immunopeptidome alterations in UKE-Mel-105b cells after 24h. **(H)** Trametinib induced immunopeptidome alterations in UKE-Mel-105b cells after 72h. **(I)** Immunopeptidomic alterations of the Ma-Mel-63a upon vemurafenib+cobimetinib treatment for 24h. **(J)** Immunopeptidomic alterations of Ma-Mel-63a upon vemurafenib+cobimetinib treatment for 48h.

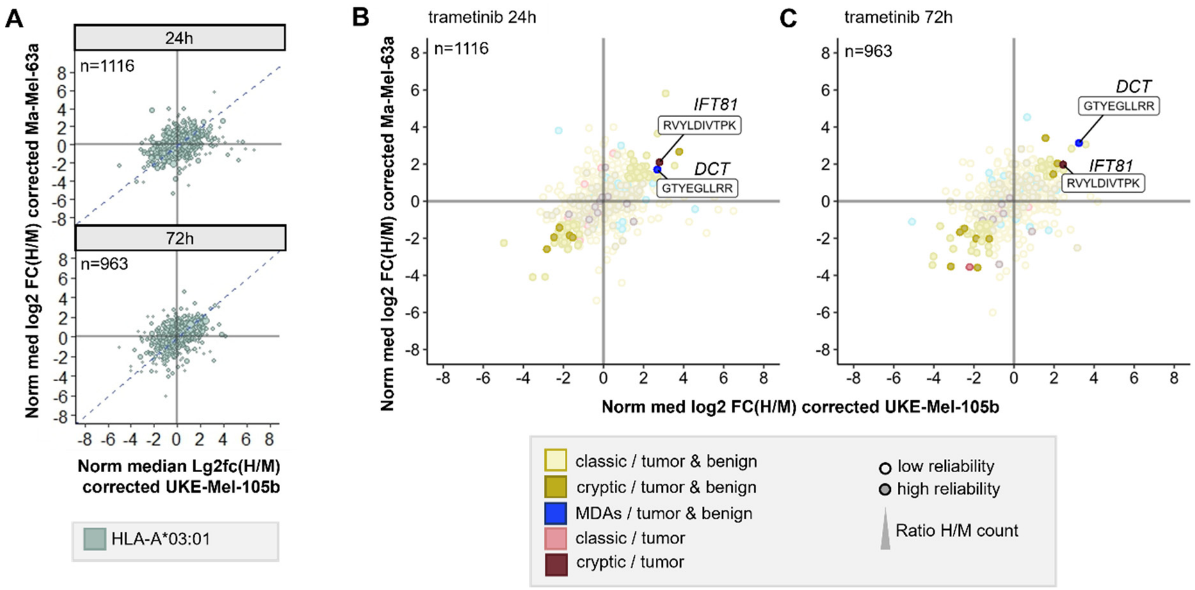


**Supplemental Figure 11 - Correlation plot of the Ma-Mel-63a and UKE-Mel-105b HLA-A*03:01 immunopeptidomic alterations upon trametinib treatment for 24h and 72h.** **(A)** Scatter plot represents correlation of shared peptides of Ma-Mel-63a and UKE-Mel-105b cells. **(B)** The cryptic peptide RVYLDIVTPK and the peptide GTYEGLLRR were upregulated in both cell lines upon trametinib treatment for 24h. **(C)** This observation could be reproduced upon 72h treatment.

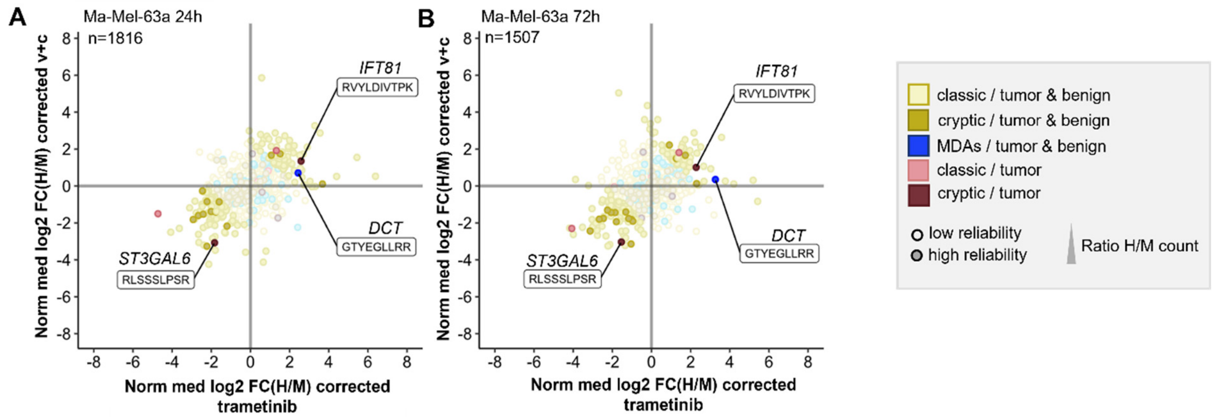


**Supplemental Figure 12 - Correlation plot of Ma-Mel-63a immunopeptidomic changes upon MAPK signaling inhibition with trametinib for 24h and 72h versus cobimetinib+vemurafenib (v+c) for 24h and 48h. (A)** The cryptic peptide RLSSSLPSR derived from *ST3GAL6* was reproducibly downregulated, while the cryptic peptide RVYLDIVTPK derived from *IFT81,* as well as the differentiation antigen GTYEGLLRR showed enhanced presentation after 24h. **(B)** This observation could be reproduced upon 72h treatment.

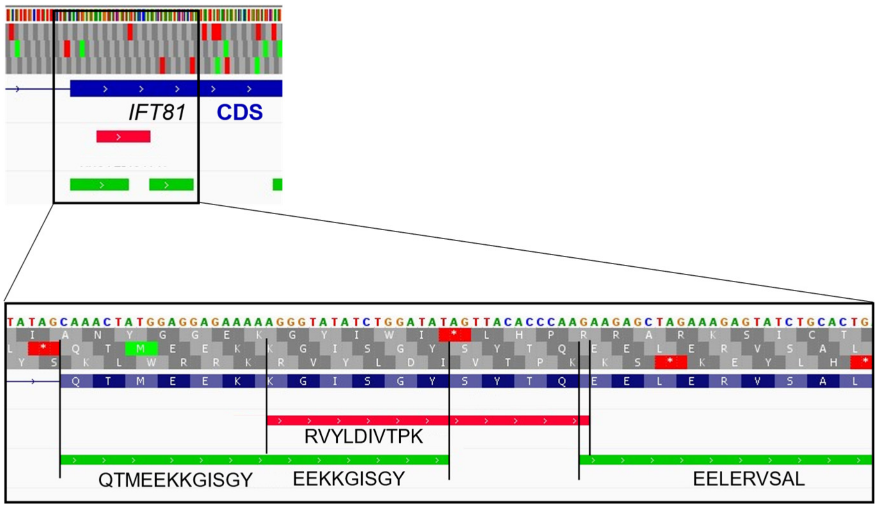


**Supplemental Figure 13 - The peptide RVYLDIVTPK was annotated as an off-frame peptide of the coding region of *IFT81*.** In contrast, classic peptides (QTMEEKKGISGY, EEKKGISGY, EELERVSAL) also derive out the coding region of *IFT81* but are in-frame. Gene locations were assessed with Integrative Genomics Viewer (1) (chr12:110190918-110190988).

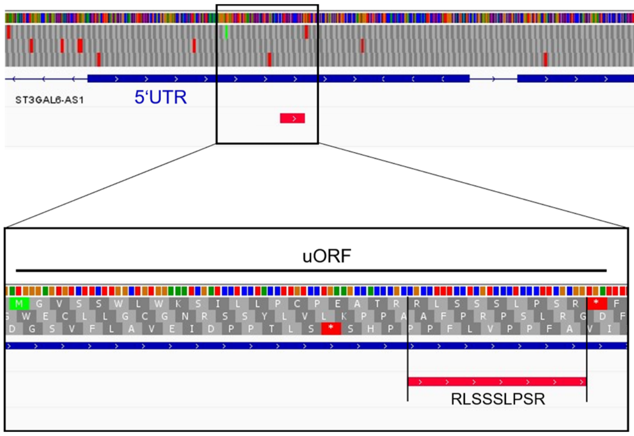


**Supplemental Figure 14 - The peptide RLSSSLPSR was annotated to an upstream open reading frame in the 5'UTR of the gene *ST3GAL6*.** Translation of the peptide starts with an CTG start codon and ends at a stop codon. Gene locations were assessed with Integrative Genomics Viewer (1) (chr3:98732425-98732492).

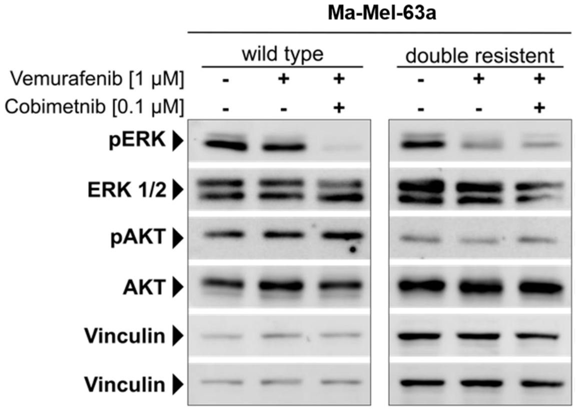


**Supplemental Figure 15 - Western Blot analyses of Ma-Mel-63a wild type cells and 2 µM Vemurafenib and 0.2 µM Cobimetinib double resistant cells, after 24h treatment with 1 µM Vemurafenib or 1 µM Vemurafenib and 0.1 µM Cobimetinib.** Inhibition of MAPK-signaling was confirmed by depletion of phospho ERK (pERK), phosphorylation at TEY motif. Additionally, the phosphorylation of AKT was investigated, as PI3K/AKT signaling is a key alternative pathway activated in BRAF and MEK inhibitor resistant cells. Vinculin was used as loading control.

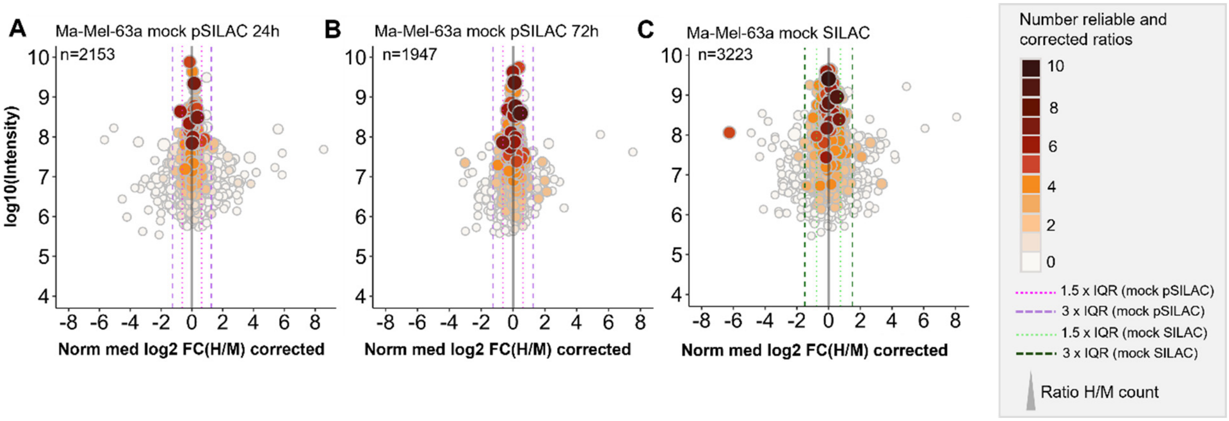


**Supplemental Figure 16 - Immunopeptidomic alterations upon pulsed-SILAC and SILAC under mock conditions.** Both labeling approaches lead to reliable quantification using our optimized data analysis strategy. **(A)** Immunopeptidome analysis after a 24-hour pulse of isotope labeling. **(B)** Immunopeptidome analysis after a 72-hour pulse of isotope labeling. **(C)** Immunopeptidome analysis of completely isotopically marked cells (SILAC)

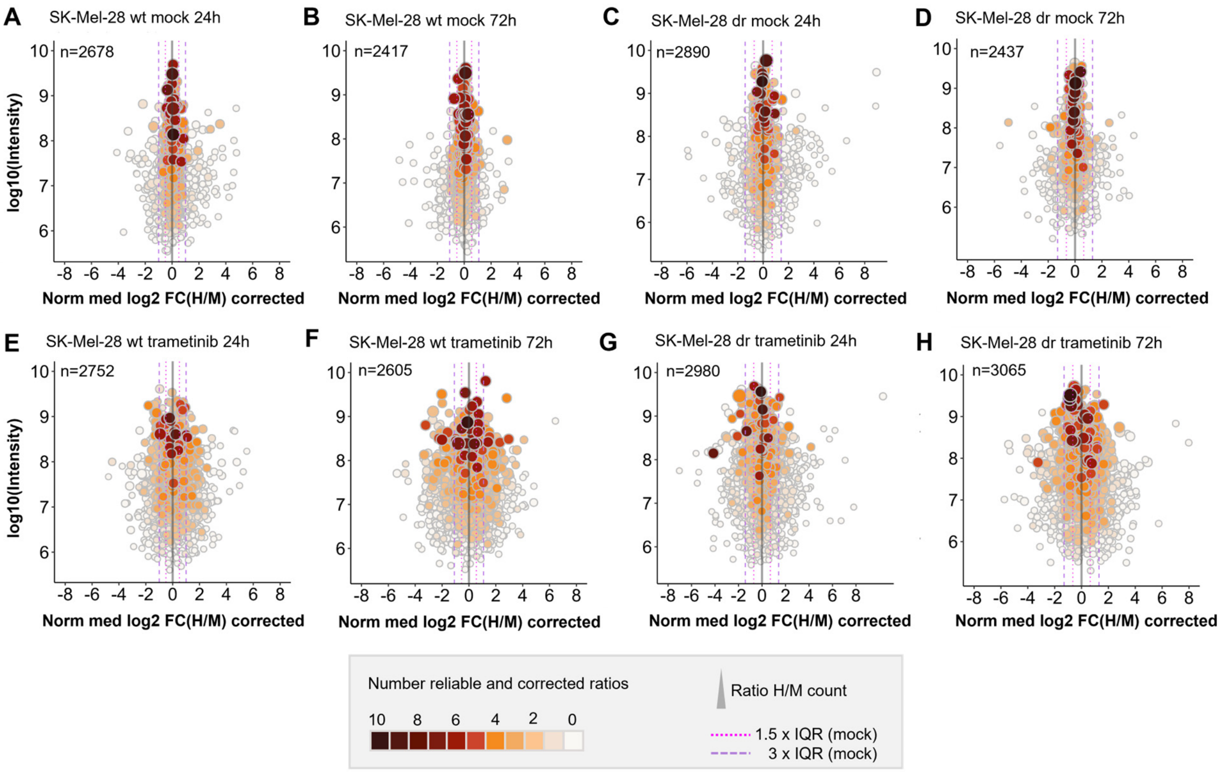


**Supplemental Figure 17 - Global immunopeptidome alterations of all tested conditions and cell lines after conducting pSILAC. (A)** Alterations of the SK-Mel-28 wt immunopeptidome upon 24h under mock conditions. **(B)** Alterations of the immunopeptidome of SK-Mel-28 wt cells upon 72h under mock conditions. **(C)** Alterations of the SK-Mel-28 dr immunopeptidome upon 24h under mock conditions. **(D)** Alterations of the immunopeptidome of SK-Mel-28 dr cells upon 72h under mock conditions. **(E)** Immunopeptidomic alterations of SK-Mel-28 wt cells upon vemurafenib+cobimetinib treatment for 24h. **(F)** Immunopeptidomic alterations of SK-Mel-28 wt cells upon vemurafenib+cobimetinib treatment for 72h. **(G)** Vemurafenib+cobimetinib induced immunopeptidome alterations in SK-Mel-28 dr cells after 24h. **(H)** Vemurafenib+cobimetinib induced immunopeptidome alterations in SK-Mel-28 rr cells after 72h.

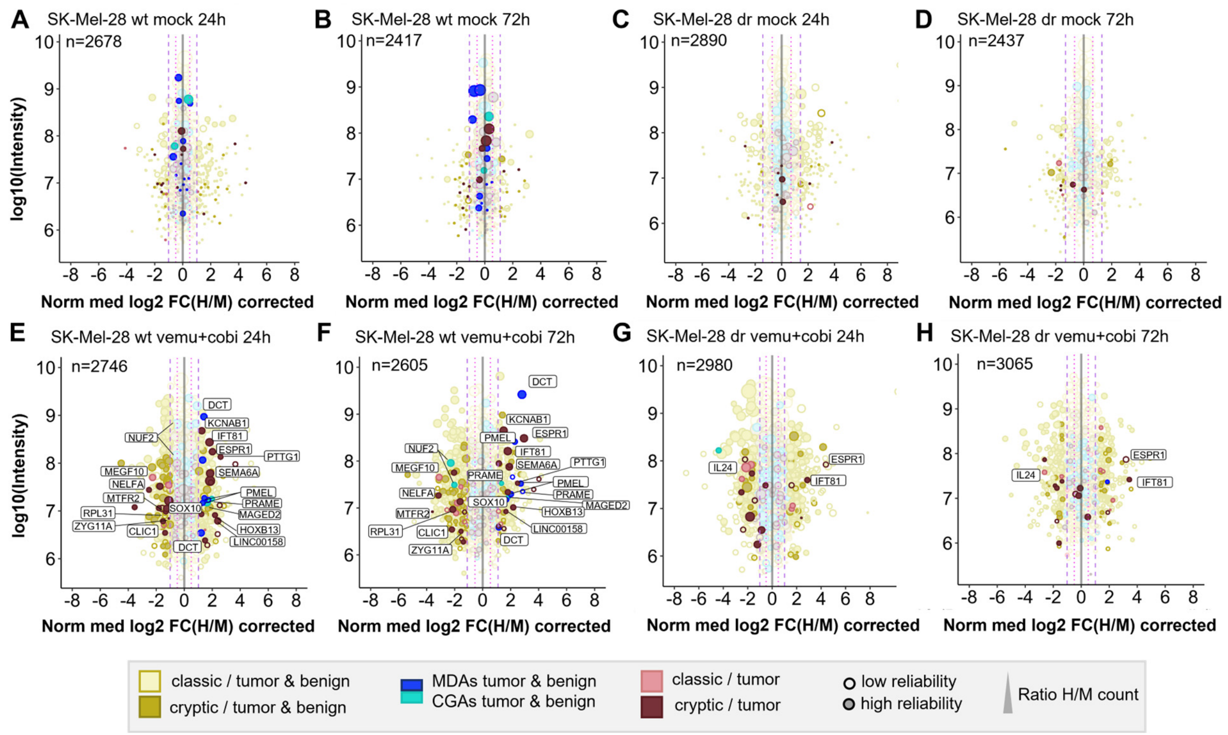


**Supplemental Figure 18 - Quantitative changes of the immunopeptide repertoires of SK-Mel-28 wt and double resistant (dr) cells under mock conditions or upon combined vemurafenib/cobimetinib treatment.** Peptides that were reliably altered in presentation upon both time points were labeled. Mock experiments were performed to estimate cell line-specific background changes occurring during labeling intervals without treatment. **(A)** Mock experiments of SK-Mel-28 wt after 24h. **(B)** Mock experiments of SK-Mel-28 wt after 72h. **(C)** Mock experiments of SK-Mel-28 dr after 24h. **(D)** Mock experiments of SK-Mel-28 dr after 72h. **(E)** SK-Mel-28 wt cells were treated with 2µM vemurafenib and 200 nM cobimetinib for 24h. **(F)** SK-Mel-28 wt cells were treated with 2µM vemurafenib and 200 nM cobimetinib for 72h. **(G)** SK-Mel-28 dr cells were treated with 2µM vemurafenib and 200 nM cobimetinib for 24h. **(H)** SK-Mel-28 dr cells were treated with 2µM vemurafenib and 200 nM cobimetinib for 72h. Immunopeptidomes were globally altered upon treatment. Presentation of MDAs was globally induced in SK-Mel-28 wt cells but not in SK-Mel-28 dr cells. The presentation of the putative tumor-exclusive peptide RVYLDIVTPK (*IFT81*) was induced upon treatment in SK-Mel-28 wt cells as well as in SK-Mel-28 dr cells.

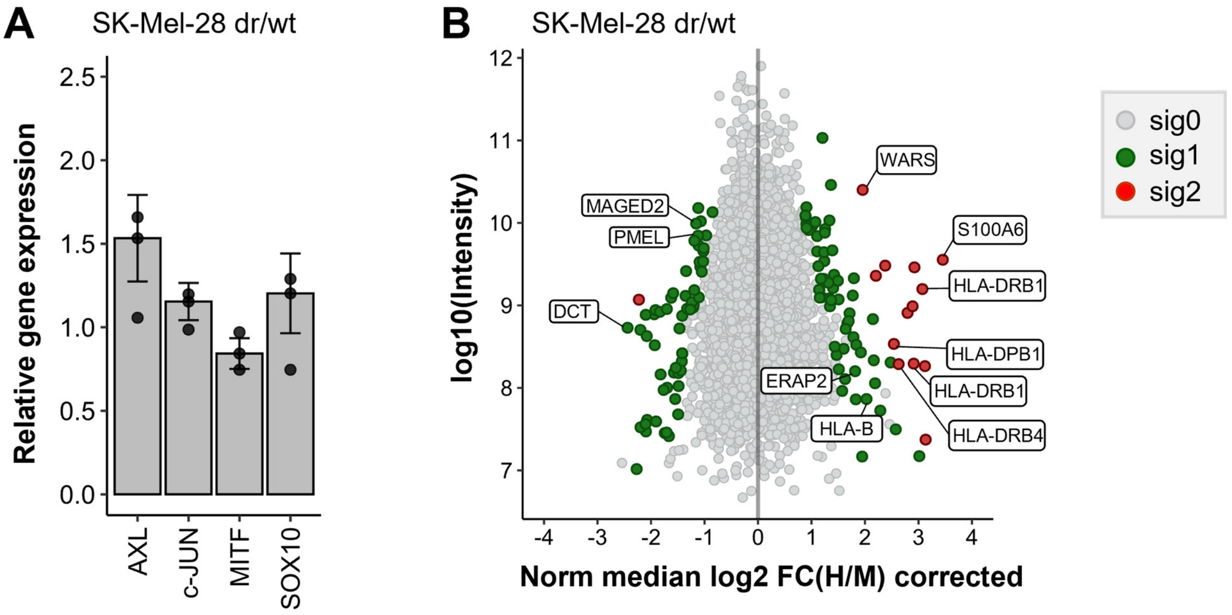


**Supplemental Figure 19 - Expression of genes that regulate or indicate melanoma phenotype switch was assessed with real time qPCR.** **(A)** Expression of melanocyte inducing transcription factor (MITF) was reduced, while expression of AXL receptor tyrosine kinase was increased in vemurafenib and cobimetinib resistant SK-Mel-28 cells in comparison to drug-sensitive wild type cells. No significant changes in expression of c-Jun and SOX10 were observed. $MITF^{low}/AXL^{high}$ expression profiles represent phenotype switching towards invasive/dedifferentiated cell states. **(B)** Quantitative proteome analysis revealed downregulation of MDAs in SK-Mel-28 dr compared to SK-Mel-28 wt cells. Apart from that, HLA-II and HLA-B expression was increased.

**Supplemental Table 1:** MaxQuant peptide table, evidence table and "extended evidence" table used for calculating H/M ratios

**Supplemental Table 2:** TvH database references

**Supplemental Table 3:** Antibodies western blot

**Supplemental Table 4:** MDAs/CGAs references

**Supplemental Table 5:** Result tables (including "extended evidence tables")

**Supplemental Table 6:** Overview of regulated putative tumor-specific cryptic peptides

**Supplemental Table 7:** Qualitative ligandome analysis of melanoctyes and keratinocytes
